# Testing Models of the Visual Word Form Area Using Combined ERP and fMRI Using the Special Properties of Chinese Characters

**DOI:** 10.1101/841817

**Authors:** Dan Xiang, Joseph Dien, Donald J. Bolger

## Abstract

The visual word form area or VWFA has been of special interest for studies of reading and dyslexia and yet there are conflicting models regarding its function. Here we put the Local Combination Detector, Lexicon, and Interactive accounts to the test, using a combination of event-related potentials and functional magnetic resonance imaging. We do so using both pseudoword and reversed radical false word manipulations with Chinese characters, making use of its special properties. We recorded event-related potentials with 68 channels while twenty native Chinese speakers were making rhyme and meaning judgments on single Chinese characters and BOLD signals were collected in a 3T magnet using multi-plane EPI with a further fifteen native Chinese speakers. The word N170 showed a prolongation for reversed radical false characters while the VWFA also showed an effect of reversal, albeit only for pseudocharacters. Furthermore, an N450 rhyming effect was observed in the phonological task compared to the semantic task, but only via an interaction with reversal. The source analysis of the N450 co-registered with a Supplementary Motor Area activation. The combination of these observations suggests that the ventral orthographic pathway is partially order insensitive and that full phonological encoding occurs relatively late, supporting and expanding a model of dyslexia. Overall, they best support a version of the Lexicon account of the VWFA.

## Introduction

The role of orthography versus phonology is one of the core research questions in how we recognize the printed word. Dueling models (Coltheart, Rastle, Perry, Langdon, & Ziegler, 2001; Plaut, McClelland, Seidenberg, & Patterson, 1996; Seidenberg & McClelland, 1989) clash, among other things, over the extent to which visual word recognition is reliant on an intermediate phonetic encoding and to what extent an orthographic code might proceed directly to semantic access. This question has important implications for literacy education (Ehri, Nunes, Stahl, & Willows, 2001; Keith E. Stanovich & Stanovich, 1995; Wyse, 2000; Wyse & Styles, 2007) and reading disorders (Castles & Coltheart, 1993; Pugh et al., 2001; K. E. Stanovich, Siegel, & Gottardo, 1997).

An area of central interest for reading researchers is the visual word form area or VWFA, which is located in the left ventral occipital-temporal cortex (OTC) in the mid-fusiform gyrus (Cohen et al., 2000, 2002; B. D. McCandliss, Cohen, & Dehaene, 2003). Growing evidence shows that the VWFA responds to words more than to other visual stimuli, such as checkerboards (Cohen et al., 2002), scrambled visual stimuli, and line drawings of objects (Baker et al., 2007; Szwed et al., 2011). Also, it responds to written words invariantly in regards to size, position, case, or font (Dehaene et al., 2004, 2001). More recent studies suggest that the location of the VWFA is also invariant to script type (Bai, Shi, Jiang, He, & Weng, 2011; D. J. Bolger, Perfetti, & Schneider, 2005; C. Liu et al., 2008), task manipulation (Ma et al., 2011; G. Xu, Jiang, Ma, Yang, & Weng, 2012) and sensory input modality (Reich, Szwed, Cohen, & Amedi, 2011). There is also interest in the role of the right hemisphere homologue of the VWFA or rVWFA but little has been written about it.

The key observation to be answered by any account of VWFA functioning is that it responds equally to real words and to pronounceable or legal (orthographically correct non-words, as in “hilsem”), but less strongly to false words (orthographically incorrect non-words, as in consonant strings like “twjk”). Even when no significant effect is observed between real words and false words (Tagamets, Novick, Chalmers, & Friedman, 2000), figures suggest the likelihood there was simply insufficient statistical power. Non-words are not just orthographically incorrect in the sense that they do not follow the rules of spelling, they are also largely non-pronounceable so they are also phonologically incorrect. Thus, the pattern can be summarized as: (RW=PW)>FW. The puzzle here is why pseudowords produce the same activity as real words; the simplest prediction is that a word recognition system should respond most strongly to real words. Instead pseudowords elicit equal, or sometimes even more, response than real words (Mechelli, Gorno-Tempini, & Price, 2003; Taylor, Rastle, & Davis, 2013).

The seminal VWFA theory is the Local Combination Detector or LCD hypothesis (Dehaene, Cohen, Sigman, & Vinckier, 2005; Vinckier et al., 2007), which holds that it focuses on sublexical analysis, specifically bigrams, as part of a processing stream with increasingly higher-level associations. Furthermore, these detectors are tuned to local bigrams, meaning that the two letters could be separated by something like one or perhaps more intervening letters. By this account, the VWFA responds to pseudowords because any orthographically correct letter string will by definition consist of appropriate letter sequences. Even if this letter string is not a word, the pairs of letters constituting it would still register as valid bigrams as experienced with real words and would still cause the VWFA to respond equivalently to real words.

An alternative view, which we will term the Lexicon hypothesis, suggests that the VWFA does in fact operate as a visual word form lexicon (Laurie S. Glezer, Kim, Rule, Jiang, & Riesenhuber, 2015; Laurie Schwarz Glezer, Jiang, & Riesenhuber, 2009). They argue that its response is not actually equivalent to real words and pseudowords; instead, they suggest that the fMRI signal reflects a strong response to real words by a few neurons and summed weak responses by many neurons to partial matches by pseudowords. They therefore argue that the only way to methods like repetition priming that allow one to distinguish between these two situations. It is not clear, however, why real words would not in addition elicit summed weak responses to partial matches, thus resulting in a stronger response than to pseudowords; perhaps some kind of lateral inhibition mechanism caps the overall activity.

Finally, according to the Interactive hypothesis (Price & Devlin, 2003, 2011), the VWFA mediates integrating multiple modalities including phonology to visual object recognition. In the case of word-like stimuli, as one reads, phonological and semantic areas produce predictions that feed back to the VWFA, helping guide its visual processing. Citing evidence that pseudowords sometimes produce not just equal but even more activation than real words (Mechelli et al., 2003), it explains that this is because more prediction error is being caused; furthermore, both should produce greater activation than a consonant string because it is sufficiently non-wordlike that there will be less predictive feedback by the other areas. This position is based on observations that the VWFA responds to many types of visual stimuli and manipulations. On the other hand, close inspection of the coordinates provided in at least some of these reports (Mechelli et al., 2003; Price, Wise, & Frackowiak, 1996) shows them in the adjoining posterior inferior temporal gyrus, also known as the language formulation area (Dien, Brian, Molfese, & Gold, 2013; Nielsen, 1946), rather than the VWFA per se, or including them in the region of interest (Mano et al., 2013).

Neuroimaging methods like functional magnetic resonance imaging (fMRI) provide a time-lapsed snapshot of activity in the first crucial second of activity, making it difficult to test the Interactive hypothesis in particular. Event-related methods or ERPs may provide the millisecond time-resolution required to resolve these issues. They are produced by the same aspect of neural activity that drives the hemodynamic response used by fMRI (Logothetis, Pauls, Augath, Trinath, & Oeltermann, 2001). Furthermore, we know that it takes time to reach the higher phonological and semantic regions (Barber & Kutas, 2007; Indefrey & Levelt, 2004; Sereno & Rayner, 2003), although see also (C. A. Perfetti & Tan, 1998; Charles A. Perfetti, Liu, & Tan, 2005; Pollatsek, Tan, & Rayner, 2000), so the initial ERP signal should largely be free of feedback signals.

It is widely thought that the VWFA is likely the generator for an ERP component, the word N170 (S. Brem et al., 2006; Dehaene, Cohen, Morais, & Kolinsky, 2015; Dien, 2009; Maurer & McCandliss, 2007), which is a left-lateralized negativity with a peak over the posterior scalp and peaking at about 170 ms after presentation of a visual word (Bentin, Mouchetant-Rostaing, Giard, Echallier, & Pernier, 1999; Maurer, Brem, Bucher, & Brandeis, 2005). It is to be distinguished from the face N170, which is right-lateralized and elicited by face stimuli (Bentin, Allison, Puce, Perez, & McCarthy, 1996; Bötzel, Schulze, & Stodieck, 1995). It reliably produces a stronger response to words than consonant strings and other non-orthographic stimuli (Bentin et al., 1999; B. D. McCandliss, Posner, & Givon, 1997). According to the phonological mapping hypothesis (Maurer & McCandliss, 2007; Bruce D. McCandliss & Noble, 2003), the word N170 has been trained in part by top-down phonological processing, resulting in a left-lateralization for alphabetic scripts, but is otherwise reflecting an unidentified orthographic process.

The word N170 behavior largely but not entirely tracks this behavior of the VWFA. For example, the N170 is usually described as responding equally to real words and pseudowords (Kast, Elmer, Jancke, & Meyer, 2010; Maurer, Brem, et al., 2005; Schendan, Ganis, & Kutas, 1998; Simon, Petit, Bernard, & Rebai, 2007), although smaller in one case (Maurer, Brandeis, & McCandliss, 2005). A combined ERP/fMRI study reported marked divergences (Silvia Brem et al., 2009), which were ascribed to fMRI being more sensitive to prolonged tonic activity and top-down influences whereas ERPs are more sensitive to the initial phasic burst of activity, which is exactly why it would be helpful for testing Interactive hypothesis (Carreiras, Armstrong, Perea, & Frost, 2014). The strongest evidence comes from intracranial reports of a strong N170-like response to word-like stimuli recorded in the vicinity of the VWFA (Allison, McCarthy, Nobre, Puce, & Belger, 1994; Mainy et al., 2008; Nobre, Allison, & McCarthy, 1994).

Thus far, a definitive test of these models has been impeded by the complexities of stimulus generation in alphabetic languages. A possible way forward is presented by the Chinese script, whose characters are comprised of radicals. As in English letters, these radicals are separate subunits that provide distinct information and may be recombined to produce new morphemes (Hsiao & Shillcock, 2006; Charles A. Perfetti et al., 2005; Siok & Fletcher, 2001). In Chinese orthography, the dominant structure is phonograms, dual-radical characters, among which about 70% are a left–right structure, with a semantic radical on the left and a phonetic radical on the right (“SP phonograms”), providing cues for semantics and phonology, respectively (Ho, Ng, & Ng, 2003). Radicals may provide phonetic and semantic cues, but researchers disagree on whether they do so to a lesser (Pollatsek et al., 2000; Tan, Laird, Li, & Fox, 2005) or greater degree (Lee, Huang, Kuo, Tsai, & Tzeng, 2010). It is unknown whether the same might be true for false characters.

The VWFA appears to mediate analysis of Chinese radicals, just as it does alphabetic letters. Meta-analyses (D. J. Bolger et al., 2005; Tan et al., 2005; Wu, Ho, & Chen, 2012; Zhao, Fan, Liu, Wang, & Yang, 2017) indicate that the VWFA, as in alphabetic languages, is robustly activated by Chinese characters, although with greater involvement of the right hemisphere homologue, perhaps due to the greater configural aspect of Chinese characters (Y. Liu & Perfetti, 2003; Tan et al., 2000) or perhaps per the phonological mapping hypothesis (Bruce D. McCandliss & Noble, 2003; Mei et al., 2013). Just as pure alexic English readers, who have damage in the vicinity of the VWFA, engage in letter-by-letter reading (Gaillard et al., 2006), pure alexic Chinese readers engage in radical-by-radical reading (Wengang & Butterworth, 1998).

The advantage of the Chinese script is that it provides a simplified system in which to manipulate bigram and lexical status. To generate a set of pseudowords matched for sublexical characteristics, one simply re-pairs the radicals such that they do not form a real word. To generate the equivalent of an English consonant string, again matched for sublexical characteristics, one simply reverses the order of the semantic and phonetic radicals to form an orthographically illegal, unpronounceable false character, although some radicals can serve in either role (Su, Mak, Cheung, & Law, 2012). Unlike in English, these two manipulations are orthogonal to each other, allowing one to readily examine the effects of bigram association strength and bigram order, and with many more potential two-subunit stimuli than in English. We suggest that a study of Chinese phonogram recognition with both a pseudocharacter and a reversal manipulation could provide a strong test of these three models.

The reversability of Chinese radicals also raises an issue thus far not addressed in the literature. At least one study of the VWFA (Dehaene et al., 2004) reported the surprising finding that under subliminal repetitive priming conditions, the VWFA did not respond to letter order scrambling. This observation is consistent with cognitive studies of orthographic processing, which has demonstrated a certain amount of order-insensitivity (Grainger, 2008). This has been most vividly demonstrated with the so-called Cambridge effect (Velan & Frost, 2007), which is that English readers are surprisingly able to read words with their internal letter order scrambled; the suggestion here being that another part of the reading system, such as the phonology pathway, could mediate order sensitivity (Frankish & Turner, 2007). The LCD model accounts for this to some degree by invoking the use of local bigrams, in that they are posited to be flexible about having 1-2 intervening characters, so the same A-C bigram detector could respond to both “ACT” and “ATC.” What is unclear is whether it could respond to an inversion of order, as in “CAT.” While order sensitivity seems to be implied in the description of the model (Dehaene et al., 2005, p. 337), it doesn’t seem to be a core feature. Likewise, the Lexicon and Interactive models could accommodate such order insensitivity depending on the details of the orthographic analysis. With Chinese phonograms, this distinction is readily tested simply by contrasting reversed radical false characters made from real characters (RF) with those made from pseudocharacters (PF).

The LCD hypothesis would predict that pseudocharacters would elicit less of a response than real characters. Since Chinese SP phonograms consist of only two radicals, pseudocharacters by definition would be invalid bigrams with two radicals that are not normally found together in a character. Thus, the predicted pattern would be: RW>(PW=RF=PF) if order-sensitive and (RW=RF)>(PW=PF) if order-insensitive.

The Lexicon hypothesis would predict that, just as with English words, the activity from summed partial matches should equal the activity from a full match of a real character. Presumably the same mechanism that results in equivalent activations for real and pseudo words in English would operate for Chinese characters. False characters, on the other hand, should result in minimal partial matches as the radicals appear in the wrong positions, if order-sensitive. Thus, the predicted pattern would be: (RW=PW)>(RF=PF) if order-sensitive and (RW=PW=RF=PF) if order-insensitive.

The Interactive hypothesis, which for Chinese is very consistent with the Lexical Constituency model (Charles A. Perfetti et al., 2005; Charles A. Perfetti, Tan, & Tan, 1999), would predict that pseudocharacters would elicit a greater response since there would be a larger prediction error from the phonology and semantic areas. Furthermore, if the VWFA is order-sensitive, false characters should produce less activity since it would be immediately apparent that they were not real characters, short circuiting the feedback from phonology and semantic areas. Thus, the predicted pattern would be: PW>RW>(RF=PF) if order-sensitive and (PW=PF)>(RW=RF) if order-insensitive.

One study (Kao, Chen, & Chen, 2010) has reported more activity to real than reversed characters in the posterior portion of the VWFA and another study (C. Liu et al., 2008) appears to have found more activity to pseudocharacters than real characters in the VWFA, but only two neuroimaging studies have examined Chinese characters with both the pseudocharacter and the reversal manipulations (Yang, Wang, Shu, & Zevin, 2011, 2012). While the results were intriguing, they were insufficient to test the present hypotheses. The second study appears to find more activity in the VWFA to phonogram pseudowords than to real words for a lexical decision task and comparable for a passive task (detect symbols). The first problem is that although they report a significant task by stimulus interaction, they did not provide pairwise contrasts so it is not known which were significantly different. The second problem is that they only had reversed pseudocharacters as false characters. The third is that the active task confounded the stimulus types of interest (e.g. real versus non-real), potentially contaminating these effects with decision-processes, by causing the participants to consider the pseudocharacters for a longer time.

As for ERP studies, for studies of native Chinese-speakers reading Chinese characters, results have thus far followed the Lexicon hypothesis (RW=PW)>FW pattern. The left N170 is reported to be equal for both real characters and pseudocharacters (Lin et al., 2011; Xue, Jiang, Chen, & Dong, 2008; Yum, Law, Su, Lau, & Mo, 2014; Zhou & Yan, 2018), with one exception (Lu, Tang, Zhou, & Yu, 2011). Furthermore, reversing the radicals of pseudocharacters to produce a false character has resulted in a smaller N170 (Lin et al., 2011; Yum, Su, & Law, 2015). An exception (Chung, Tong, & McBride-Chang, 2012) apparently did not include the peak N170 channels in the analysis.

As a final element in this study, we will use the N450 as a reliable method for detecting the use of phonology. The N450 is a strong negativity to non-rhyming versus rhyming stimuli, peaking in the 400-500 ms range over Cz-Pz (M. D. Rugg, 1984a, 1984b). This effect has also been demonstrated in children (Coch, Grossi, Skendzel, & Neville, 2005; Grossi, Coch, Neville, Coffey-Corina, & Holcomb, 2001) and has been shown to be diminished in dyslexia (Ackerman, Dykman, & Oglesby, 1994). Since the task was rhyme detection in these studies, it is likely that the effect was contaminated by a P300 (Pz peak) and so the actual N450 peak channel would be more central. The N450 effect is strongly right-lateralized for orthographically dissimilar stimuli (e.g., MAKE-ACHE) and midline for similar stimuli (e.g., PLEA-FLEA) for reasons that are currently unclear (Michael D. Rugg & Barrett, 1987). This effect is also found for pronounceable pseudowords (Ackerman et al., 1994; M. D. Rugg, 1984a), single letters (Coch, Hart, & Mitra, 2008), auditory words (Mitra & Coch, 2019; Perez-Abalo, Rodriguez, Bobes, Gutierrez, & Valdes-Sosa, 1994), and even nameable pictures (Barrett & Rugg, 1990; Coch, 2018). Its generator site is unknown and localizing it will be a secondary goal for this study.

The N450 effect has also been reported with Chinese characters (Chen, Lee, Kuo, Hung, & Cheng, 2010; Valdes-Sosa et al., 1993). In a prior study (Y. Liu, Perfetti, & Hart, 2003), an N450 is also very visible in a homophone task but not a meaning task. The former study displayed the right parietal N450 topography while the latter was mostly the bilateral central N450 topography. No enough information was provided regarding the latter study’s stimuli to determine if this difference between studies was mediated by orthographic similarity of the character pairs.

In the present study, two separate samples of native Chinese speakers, one for EEG and one for fMRI, scanned a series of Chinese characters for targets. The two tasks required full character recognition (semantic judgment and rhyming judgment) and the rare target stimuli were not analyzed to avoid decision-making confounds. Non-target stimuli consisted of real characters, pseudocharacters, reversed real characters, reversed pseudocharacters, and false stroke non-characters. We also added a 250 ms SOA backward mask to reduce the influence of post-lexical processing.

### 2.1. ERP experiment material and methods

#### 2.1.1. Participants

Twenty volunteers participated in the EEG experiment in exchange for honoraria (9 male, 11 female; mean 23.3 years old, range 19-31), following a protocol approved by the University of Maryland Institutional Review Board; written informed consent was obtained from all participants. There were no conflicts of interest. All had normal or corrected-to-normal vision, were right handed, and were native Chinese mandarin speakers from the mainland and hence used to simplified characters. None reported any history of neurological, psychiatric, or learning disorders.

#### 2.1.2. Stimuli

A total of 208 simplified Chinese SP phonograms, with a semantic radical on the left and a pronounceable phonetic radical on the right, half regular and half irregular, were selected from the Chinese Single Character Word Database (Y. Liu, Shu, & Li, 2007) at http://blclab.org/pyscholinguistic-norms-database/. Mean and standard deviation values of the 13 main properties of the characters are presented in Table 1.

**Table 1.**
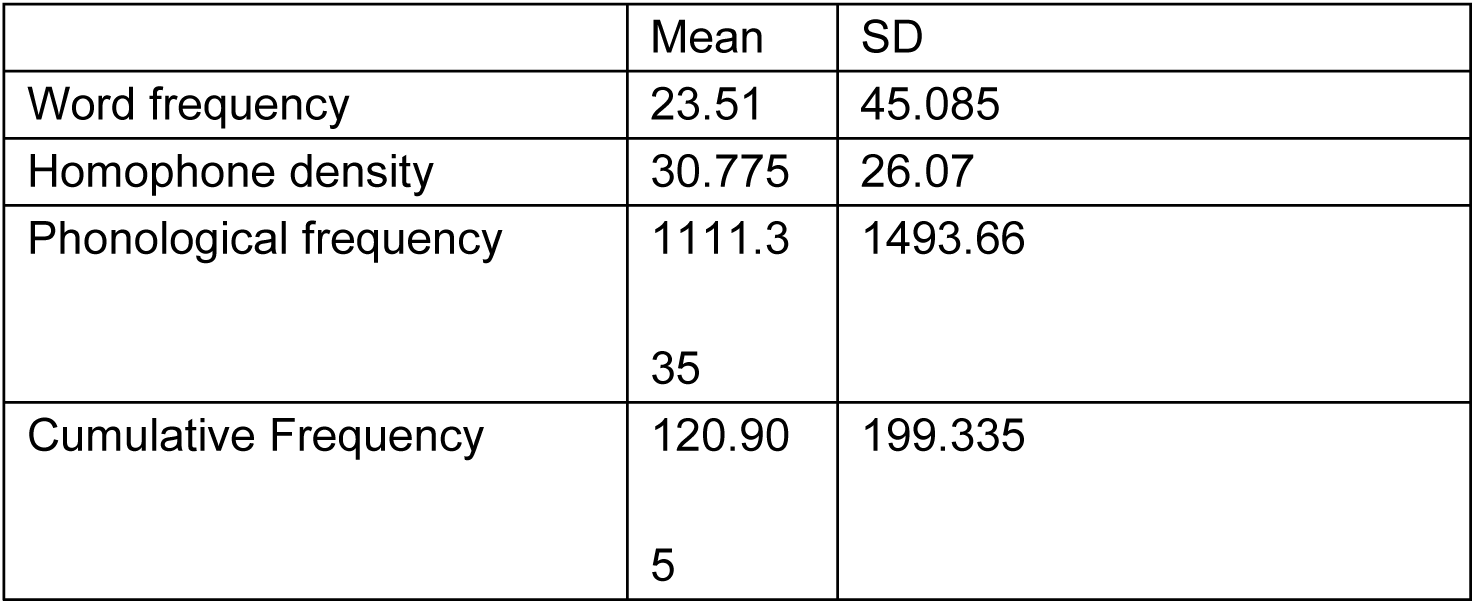
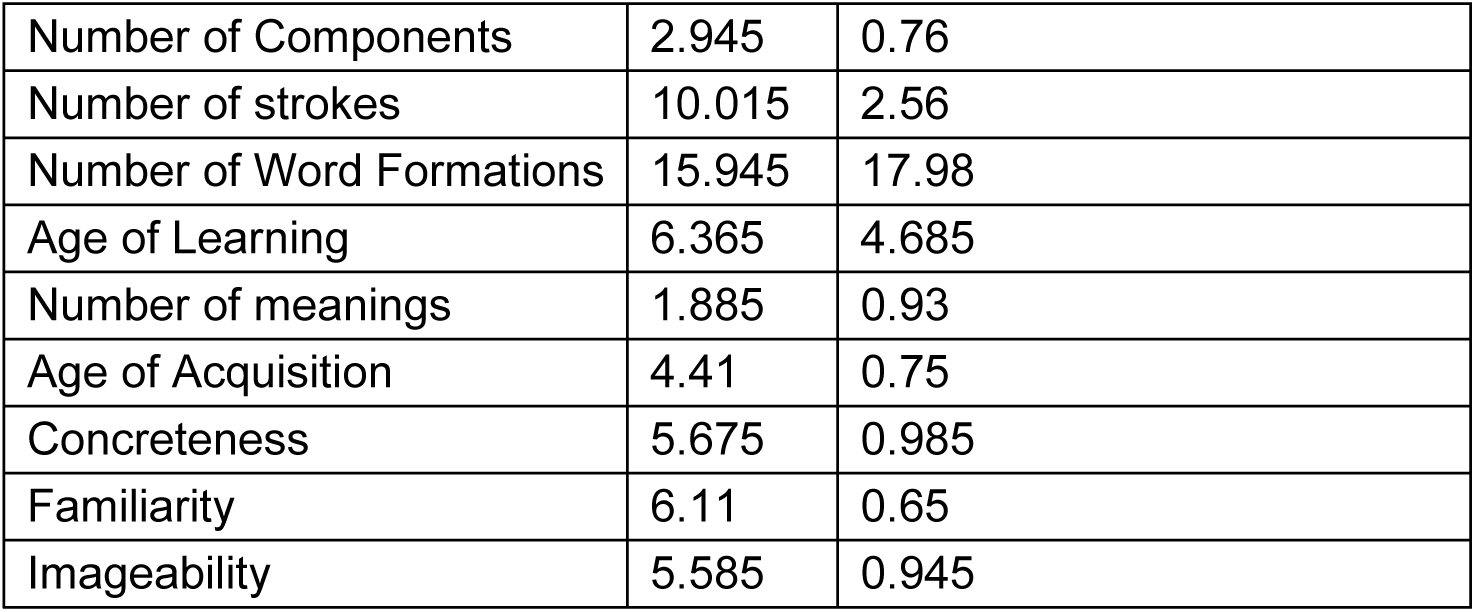
Mean, standard deviation for real characters

**Table 2.**
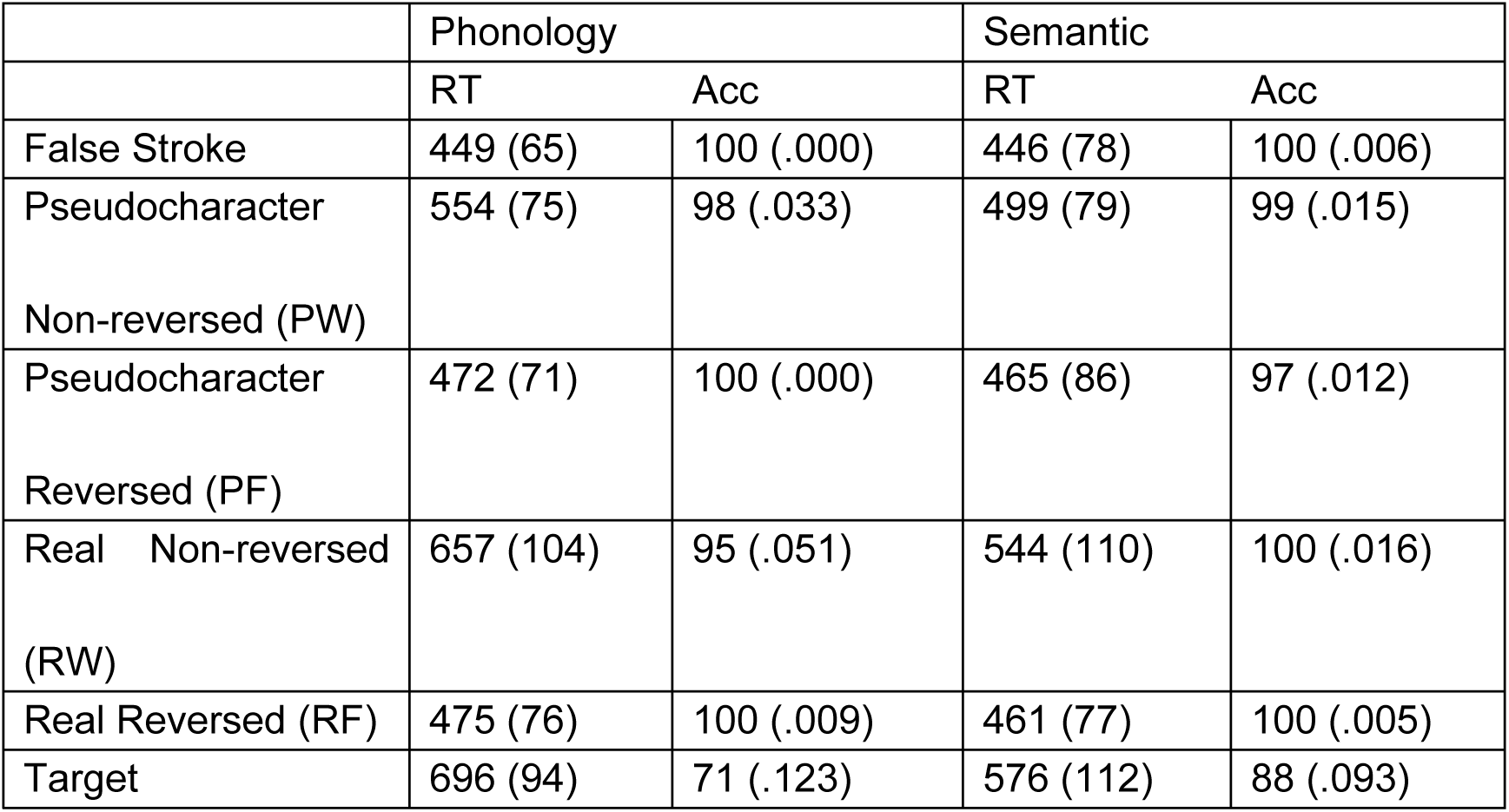
Mean of ERP Participant Median Reaction Times and Accuracy Rates. Standard deviations in parentheses.

**Table 3.**
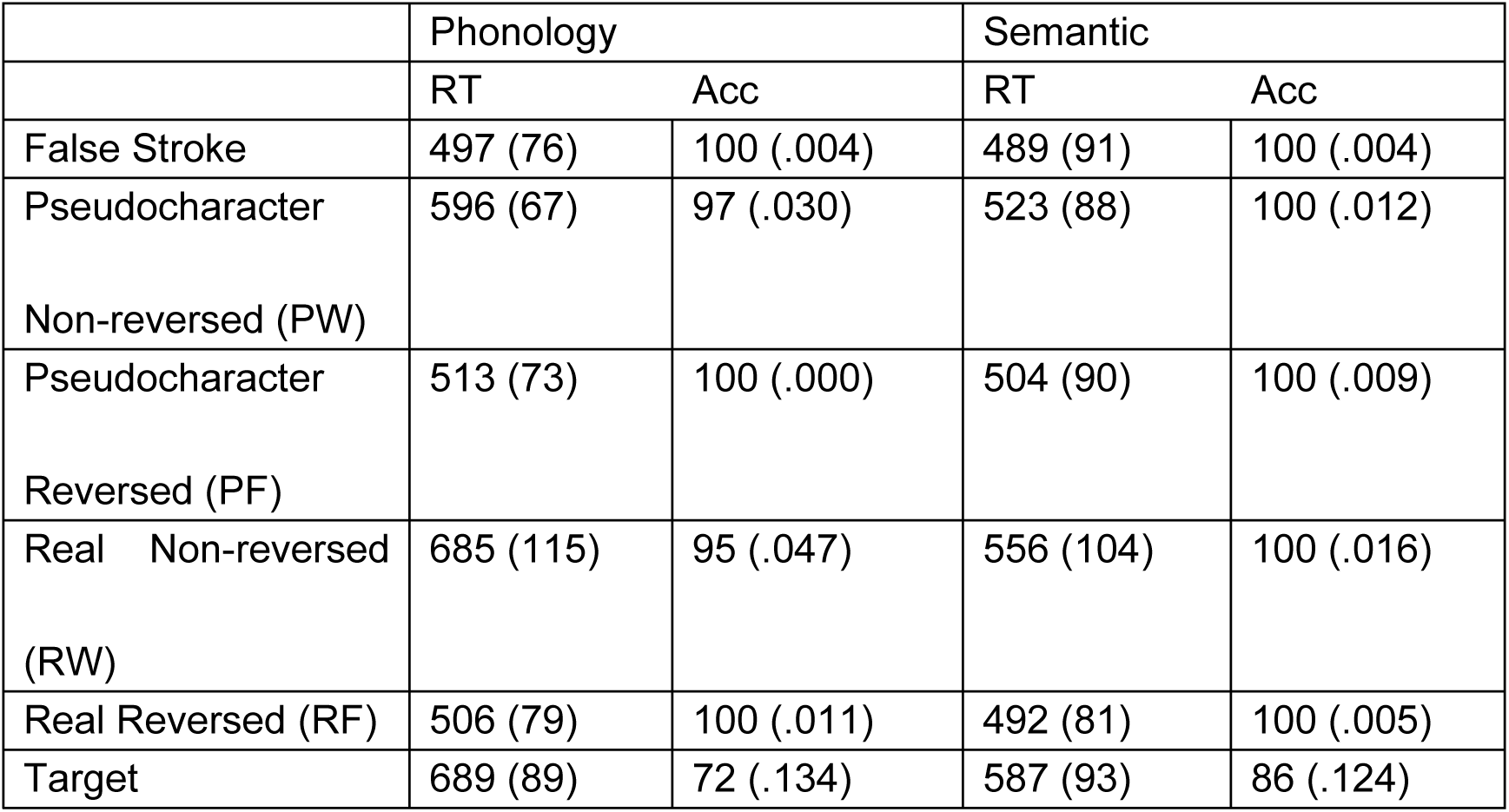
Mean of fMRI Participant Median Reaction Times and Accuracy Rates. Standard deviations in parentheses.

With the phonograms as base characters, a total of 104 left/right structure pseudocharacters (PF) were created by mismatching the semantic radical and the phonetic radical randomly selected half from regular and half from irregular characters. Pseudocharacters were neither pronounceable nor meaningful at whole character level, but with orthographic legality (e.g. “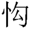”, a mismatching of semantic radical from character “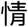” feeling /qing2/ and phonetic radical from character “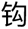” hook /gou1/; “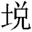”. Whether a re-pairing of a semantic radical and a phonetic radical was a pseudocharacter was decided by its availability in Xinhua Dictionary (11^th^ Edition). Further, reversed real characters (RF) and reversed pseudocharacters (PF) were constructed by switching the left/right radical position. A total of 208 false stoke combinations were also built by randomly-combining legal strokes in a square shape configuration, each with a same number of strokes as its real character counterpart. An additional 48 SP phonograms were used as target stimuli, half phonological and half semantic targets, also regularity was balanced. A pattern backward mask “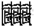” was used.

#### 2.1.3. Procedure

Experiment sessions were run using E-Prime 2.0.10.353 (Psychology Software Tools Inc., Pittsburgh PA). The tasks were rhyming and meaning judgment. For the rhyming judgment, participants determined if the presented stimulus rhymed with an ‘an’ sound regardless of tone. For the meaning judgment, participants determined if the stimulus meant ‘animal’. The selection of non-target real characters excluded those rhyming with “an” sound at the radical level in the phonological task and those with a semantic clue of ‘animal’ at the radical level in the semantic task.

Participants were first asked for two practice sessions, one phonological and one semantic, with ten trials each (using stimuli not used in the experiment itself). The participant’s task was to make a decision as quickly and accurately as possible. The participants were told to ignore the backward mask. The practice sections were repeated until the participant achieved 80% correct. Four runs of 104 non-target and 12 target trials each then followed, two for each task, each lasting 7.5 min. As shown in Figure 2, stimuli were presented visually in a sequential order, where each stimulus was presented for 250 ms followed by a backward mask for 150 ms. Then, a fixation plus of 1850 ms appeared, and finally a 500 ms feedback period as to whether the response to each stimulus was correct or not. During the feedback period, the fixation turned blue if correct, red if incorrect, and teal if there was no response within 1000 ms.

**Figure 1:**
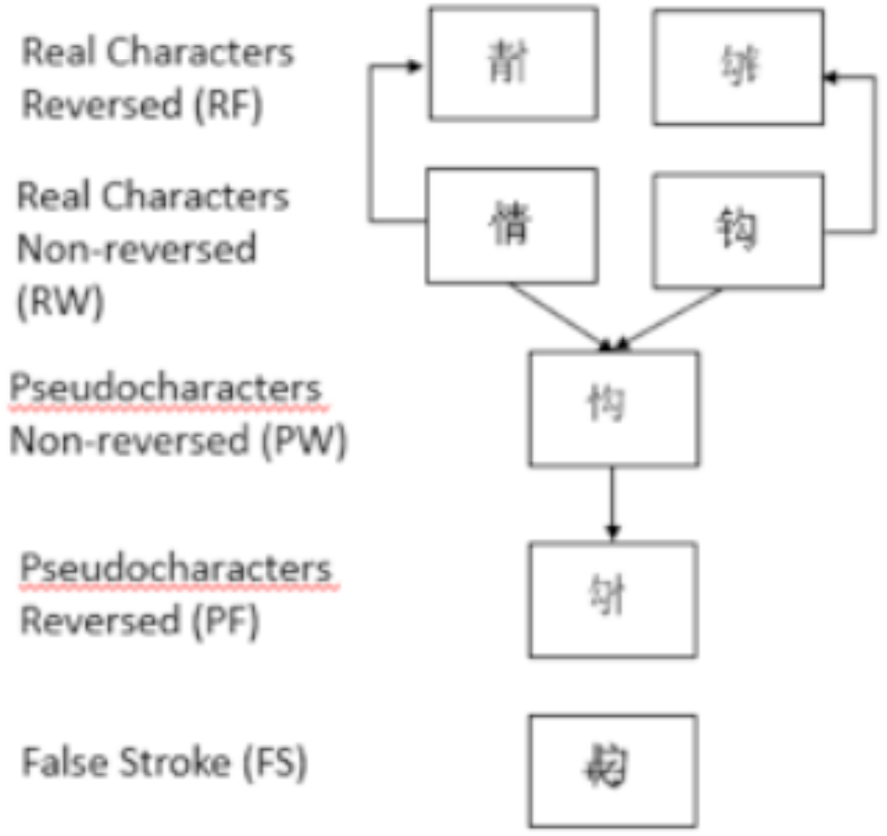
Examples of Stimuli.

**Figure 2:**
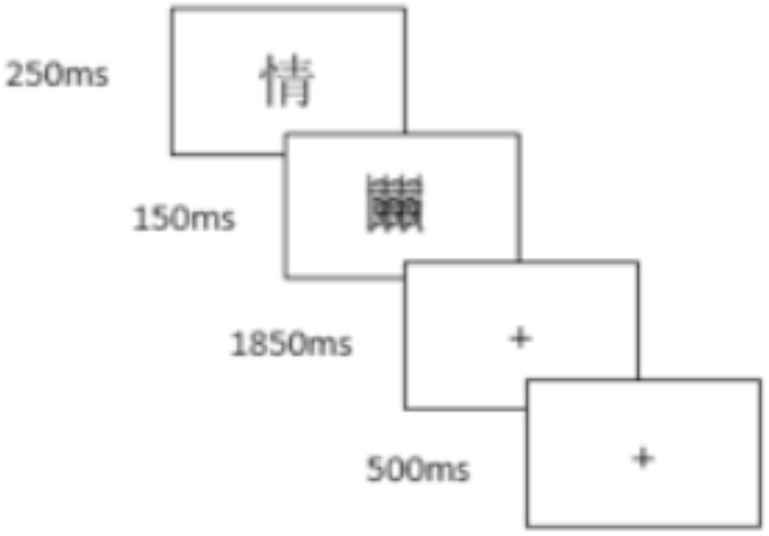
Timeline of Stimulus Presentation.

In order to maintain comparability with the fMRI data, the stimulus presentation sequence was determined in the same manner, with the inter-trial interval ranging from a minimum of 1000 ms up to several seconds by the insertion of null events (Dale, 1999). A total of four pseudorandom orders were used (randomly chosen for each run) ordered for optimal experimental efficiency using optseq2 (http://surfer.nmr.mgh.harvard.edu/optseq/). While the sequence of the trial types was determined by optseq2, the stimuli within that type was chosen randomly by E-Prime.

Furthermore, there were four stimulus lists counterbalanced between participants. Each word stimulus was rotated through the four combinations of the task and reversal manipulations, such that a participant would see a given character once during the session, with the cell determined by the list counterbalance.

The participant rested his/her chin on a chinrest 57 cm from the LED computer monitor. The stimuli were presented in black against a white background in a font of Sonti in the middle of the screen (1.8 degrees visual angle). Responses were made via button-presses, using the millisecond-accurate Black Box Toolkit button-box, and were registered with the left and right forefingers (target/non-target button counterbalanced across participants).

#### 2.1.4. Data collection

ERPs were recorded at a 1000 Hz sampling rate using a Brain Products actiCAP active electrode cap, an actiCHamp amplifier, and PyCorder 1.07. There were 64 channels on the cap plus four EOG channels. The data were referenced to a reference channel internal to the amplifier. The data were analog-filtered with a high-pass of .01 and a low pass of 200 Hz and a notch filter at 60 Hz. The impedance criterion was 17k ohms, which is the default setting for the system.

#### 2.1.5. Data analysis

The data were analyzed using the EP Toolkit (Dien, 2010) 2.72 through 2.80, available for download at https://sourceforge.net/projects/erppcatoolkit/), using the Multiple Algorithm Artifact Correction (MAAC) procedure (Dien, in preparation). First, the data were segmented from −200 to 1000 ms around the stimulus presentation. Then the segments were baseline corrected to serve as a lossless high-pass filter. Then global bad channels were identified as those whose best absolute correlation with neighboring channels falls below .4. Then the data were corrected for corneo-retinal dipole artifacts using a regression algorithm. Next, blink correction was performed via independent components analysis (Delorme & Makeig, 2004) by removing factors that correlated .9 with an automatically generated eyeblink template (Frank & Frishkoff, 2007). Then movement artifacts were removed by computing a temporal PCA on each trial and then removing factors with a maximum voltage range of over 200 µv. Then high-frequency noise was removed using blind source separation canonical correlation analysis (BSS-CCA) (De Vos et al., 2010), thus avoiding the need for a low-pass filter. Next, trialwise bad channels were identified and replaced using interpolation. Finally, data were baseline corrected and average referenced (Bertrand, Perrin, & Pernier, 1985; Dien, 1998) and down sampled to 250 Hz.

ANOVAs were carried out with three factors: Task (Phonological vs. Semantic), Reversal (radicals in normal positions vs. reversed), and Lexical (Pseudocharacter vs. Real). For the N170 analyses, there was also a Hemisphere factor (Left vs. Right). No significant effects were found for the regularity cells so they were collapsed together. The false stroke and target conditions were not of interest and hence dropped from the analysis.

There were 26 trials per cell. Only trials that were correct and had reaction times larger than or equal to 100 ms were included in the averaged waveforms and the reaction time analyses. Only trials that had reaction times larger than or equal to 100 ms were included in the accuracy analyses. Ignoring the target and the false stroke conditions, the average number of trials per condition per participant was 24.3 (range of 12 to 26 for individual participants).

Measures of the ERP components were computed based on first computing an average waveform across the channel group. Amplitude was measured as the mean voltage within the window. Latency was measured as the time of the minimum/maximum peak within the window. If there was no local peak, then it was treated as missing data. Based on the literature, the N170 was measured within a 150-200 ms window at channels centered on PO7 (O1-P5-P7-PO3-PO7) and PO8 (O2-P6-P8-PO4-PO8). The N450 was measured within a 400-500 ms window at channels centered on Cz (C1-C2-Cz-CP1-CP2-CPz-FC1-FC2-FCz).

Robust ANOVAs (denoted by “TWJt/c”) were used to test effects (Dien, 2017; Keselman, Wilcox, & Lix, 2003), as implemented in the EP Toolkit (Dien, 2010). The number of iterations used for the bootstrapping function was 49,999. The reported statistical results are the median value of 11 such replications to ensure stability. If an effect does not meet the alpha criterion for all 11 replications, then it is reported as being a borderline effect even if the median p-value meets the criterion. P-Values are rounded to the second significant digit (where available). Condition means where given are the trimmed means.

Sample-By-Sample Tests: Sample-by-sample tests were conducted using a non-parametric Wilcoxon signed rank test for a given channel. For t-maps, an additional constraint was imposed that only time points that were part of a series of significant time points would be considered significant. These tests were not subjected to multiple comparison correction and so should be considered to be descriptive follow-ups to inferential testing.

Source Analysis: Source analyses were conducted on the factors using FieldTrip’s (Oostenveld, Fries, Maris, & Schoffelen, 2011) ft_dipolefitting function, which first performed a grid search to find the best starting position, and then a non-linear fitting procedure (Lutkenhoner, 1998) to find the optimal position for an x-axis symmetric equivalent dipole pair.

Since an equivalent dipole solution only provides the source location if it was a dimensionless point, it is then necessary to determine the implied cortical source for such a solution. Using a novel approach, Matlab was used to place the dipoles into an MNI space brain model included with SPM8 (cortex_5124.surf.gii). Cones were projected along the dipole orientations with an arbitrarily chosen radius that would result in roughly about a 1 cm intersection with the cortex. The intersection point was plotted along with the fMRI clusters to allow for visual inspection of convergence. Additionally, the median coordinates of the cortical vertices that lay within the intersection cones were computed to provide a rough midpoint of the implied cortical solutions. This procedure provides two potential solutions, one at the negative voltage pole and one at the positive voltage pole. This method and its implications will be further described elsewhere (Dien, in preparation).

#### 2.1.6. Data Repository

The ERP and fMRI datasets may be downloaded from: https://umd.box.com/s/k0j460ysa4f0dz8cx9u1j2xxp77jqvhu

### 2.2. ERP experiment results

#### 2.2.1. Behavioral data

Using trimmed means, overall semantic judgments (490 ms) were faster than phonology judgments (538 ms): T_WJt_/c(1.0,17.0)=28.50, p=0.00014, MSe= 2889. Also, overall responses to pseudocharacters (497 ms) were faster than to real characters (532 ms): T_WJt_/c(1.0,17.0)=56.04, p<0.00000001, MSe= 780. Also, overall responses to reversed characters (467 ms) were faster than to non-reversed characters (562 ms): T_WJt_/c(1.0,17.0)=266.71, p<0.00000001, MSe= 1213. There was an interaction between Task and Lexical factors (T_WJt_/c(1.0,17.0)=14.13, p=0.0015, MSe= 651) such that the Lexical effect was more significant for phonology judgments (T_WJt_/c(1.0,17.0)=58.35, p<0.00000001, MSe= 797) than for semantic judgments (T_WJt_/c(1.0,17.0)=10.09, p=0.0063, MSe= 634). Furthermore, there was an interaction between Task and Reversal factors (T_WJt_/c(1.0,17.0)=42.65, p=0.000020, MSe= 1278) such that the Task effect was significant for non-reversed characters (T_WJt_/c(1.0,17.0)=52.35, p<0.00000001, MSe= 2587) but not reversed characters. Additionally, there was an interaction between Lexical and Reversal factors (T_WJt_/c(1.0,17.0)=41.06, p=0.000020, MSe= 1199) such that the Lexical effect was significant for non-reversed characters (T_WJt_/c(1.0,17.0)=59.40, p=0.000020, MSe= 1564) but not reversed characters. Finally, there was a three-way interaction between Task, Lexical, and Reversal factors: T_WJt_/c(1.0,17.0)=13.32, p=0.0039, MSe= 447. This took the form of the Task by Lexical interaction only being significant for non-reversed characters: T_WJt_/c(1.0,17.0)=20.94, p=0.00040, MSe= 715. This interaction in turn resulted from the Lexical effect being much more significant for the phonology task (T_WJt_/c(1.0,17.0)=79.19, p<0.00000001, MSe= 1152) than for the semantic task (T_WJt_/c(1.0,17.0)=14.76, p=0.0018, MSe= 1127).

The accuracy analyses yielded a similar pattern. Overall semantic judgments (100%) were more accurate than phonology judgments (98%): T_WJt_/c(1.0,17.0)=21.27, p=0.00066, MSe= 0.00033. Also, overall responses to pseudocharacters (99%) were slightly more accurate than to real characters (99%): T_WJt_/c(1.0,17.0)=5.52, p=0.029, MSe= 0.00034. Also, overall responses to reversed characters (100%) were more accurate than to non-reversed characters (98%): T_WJt_/c(1.0,17.0)=29.30, p=0.00010, MSe= 0.00039. Also, there was an interaction between Task and Lexical factors (T_WJt_/c(1.0,17.0)=10.12, p=0.0042, MSe= 0.00025) such that the Task effect was significant for real characters (T_WJt_/c(1.0,17.0)=29.92, p=0.000060, MSe= 0.00030) but not pseudocharacters. Furthermore, there was an interaction between Task and Reversal factors (T_WJt_/c(1.0,17.0)=26.38, p=0.000040, MSe= 0.00029) such that the Task effect was significant for non-reversed characters (T_WJt_/c(1.0,17.0)=29.92, p=0.000060, MSe= 0.00030) but not reversed characters. Finally, there was an interaction between Lexical and Reversal factors (T_WJt_/c(1.0,17.0)=11.46, p=0.0048, MSe= 0.00019) such that the Lexical effect was significant for non-reversed characters (T_WJt_/c(1.0,17.0)=8.57, p=0.0098, MSe= 0.00047) but not reversed characters. The three-way interaction just missed significance.

#### 2.2.2. ERP data

The bilateral N170 (Figure 3) was more delayed during the phonological task (T_WJt_/c(1.0,17.0)=5.57, p=0.033, MSe= 37.10) and for reversed characters (T_WJt_/c(1.0,17.0)=8.95, p=0.0075, MSe= 54.86). This latter effect was subject to an interaction with hemisphere (T_WJt_/c(1.0,17.0)=4.91, p=0.047, MSe= 43.47) such that it was only significant over the left hemisphere (T_WJt_/c(1.0,17.0)=9.20, p=0.0099, MSe= 73.47). According to a sample-by-sample test, these effects were mostly due to a prolongation of the N170 to the reversed characters rather than a delay. There were no significant amplitude effects for the N170.

**Figure 3:**
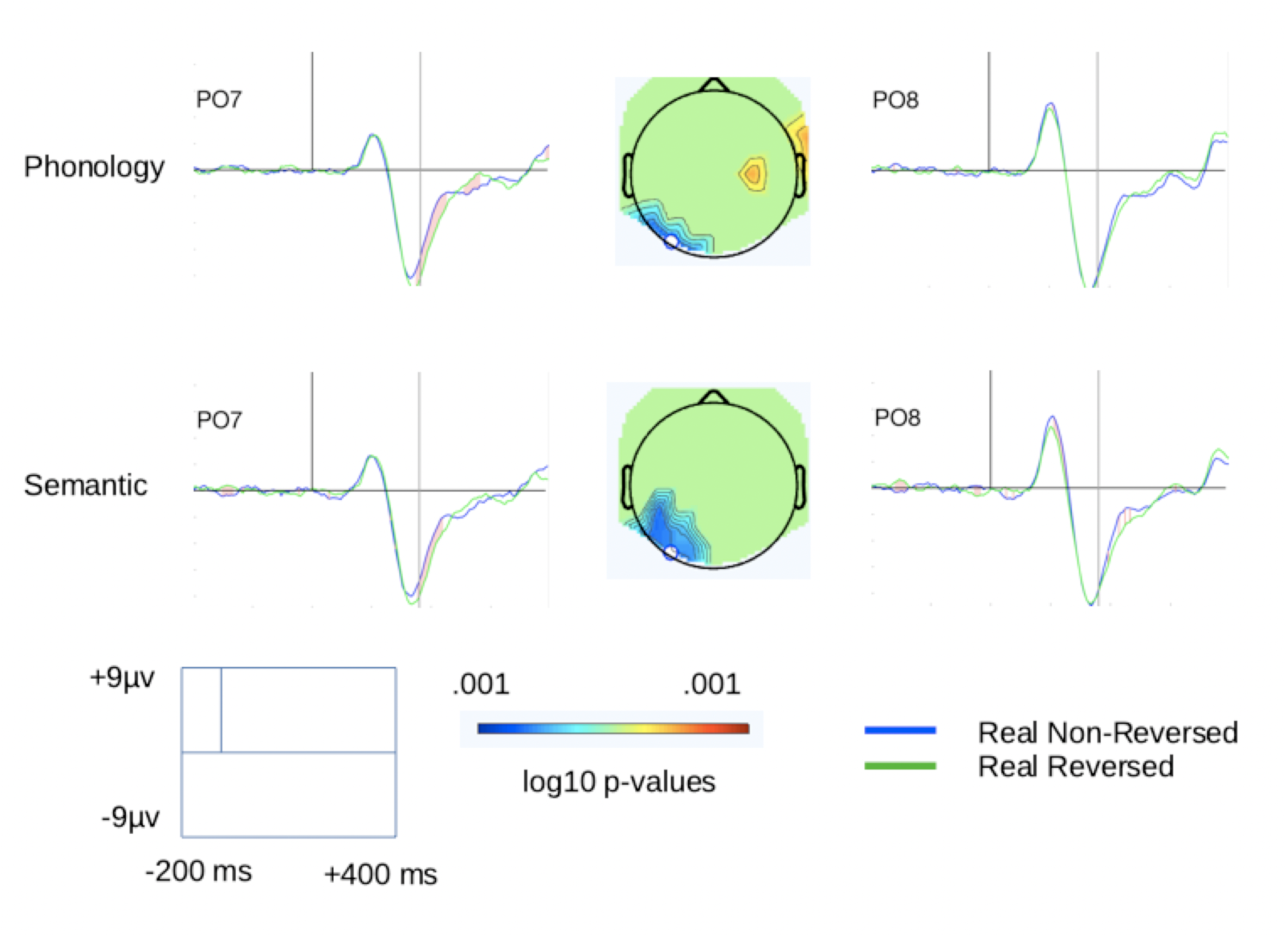
N170 Reversal Effect. Red shading shows results of sample-by-sample tests that confirm the prolongation of the N170 for reversed characters over the left hemisphere. The short black vertical line marks stimulus onset time. The long black vertical line marks 180 ms, the latency of the t-maps. The t-maps show the topography of the sample-by-sample tests. The white dot shows the location of PO7.

The N450 (Figure 4) was borderline more negative for the phonological task: T_WJt_/c(1.0,17.0)=4.42, p=0.052, MSe= 1.50. It was also more negative for the non-reversed versus the reversed characters: T_WJt_/c(1.0,17.0)=13.64, p=0.0022, MSe= 5.34. Finally, these effects interacted (T_WJt_/c(1.0,17.0)=5.55, p=0.025, MSe= 1.09) such that the Task effect was significant for non-reversed (T_WJt_/c(1.0,17.0)=12.79, p=0.0053, MSe= 0.99) but not reversed characters.

**Figure 4:**
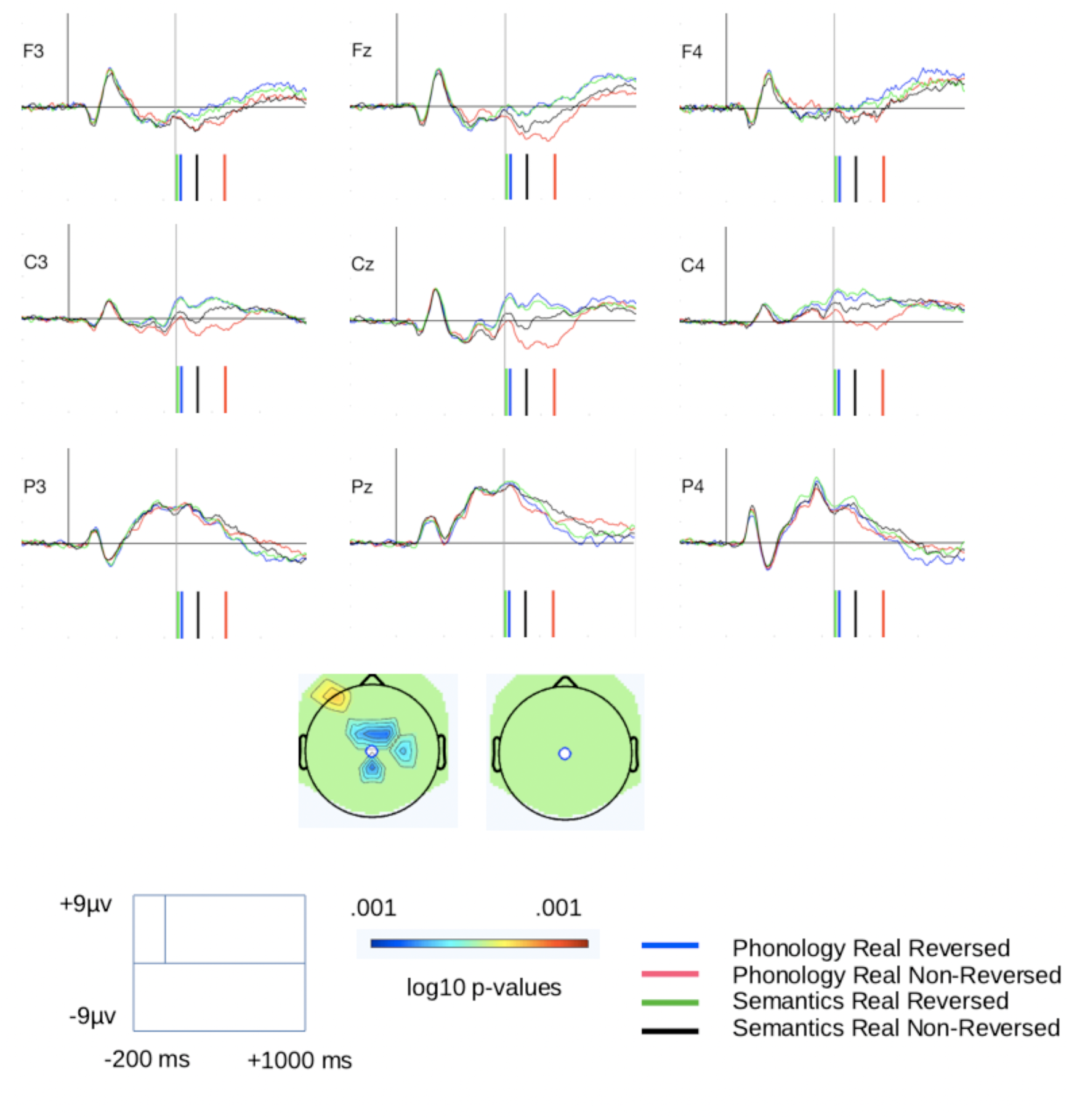
N450 Phonology Reversal Effect. The colored lines mark the reaction times. The short black vertical line marks stimulus onset time. The long black vertical line marks 450 ms, the latency of the t-maps. The t-maps show the topography of the sample-by-sample tests. The white dot shows the location of Cz.

A source analysis of the N450 effect (phonology non-reversed real character minus phonology reversed real character) at 620 ms resulted in a single dipole solution after the symmetrical dipole pair solution failed due to their converging (Figure 5). The MNI coordinates of the point equivalent dipole were [5.06 36.45 7.99], accounting for 91.52% of the variance. The implied negative cortical coordinates are [7.9 31.6 50.4] and the implied positive cortical coordinates are [5.7 37.7 −22.6].

**Figure 5:**
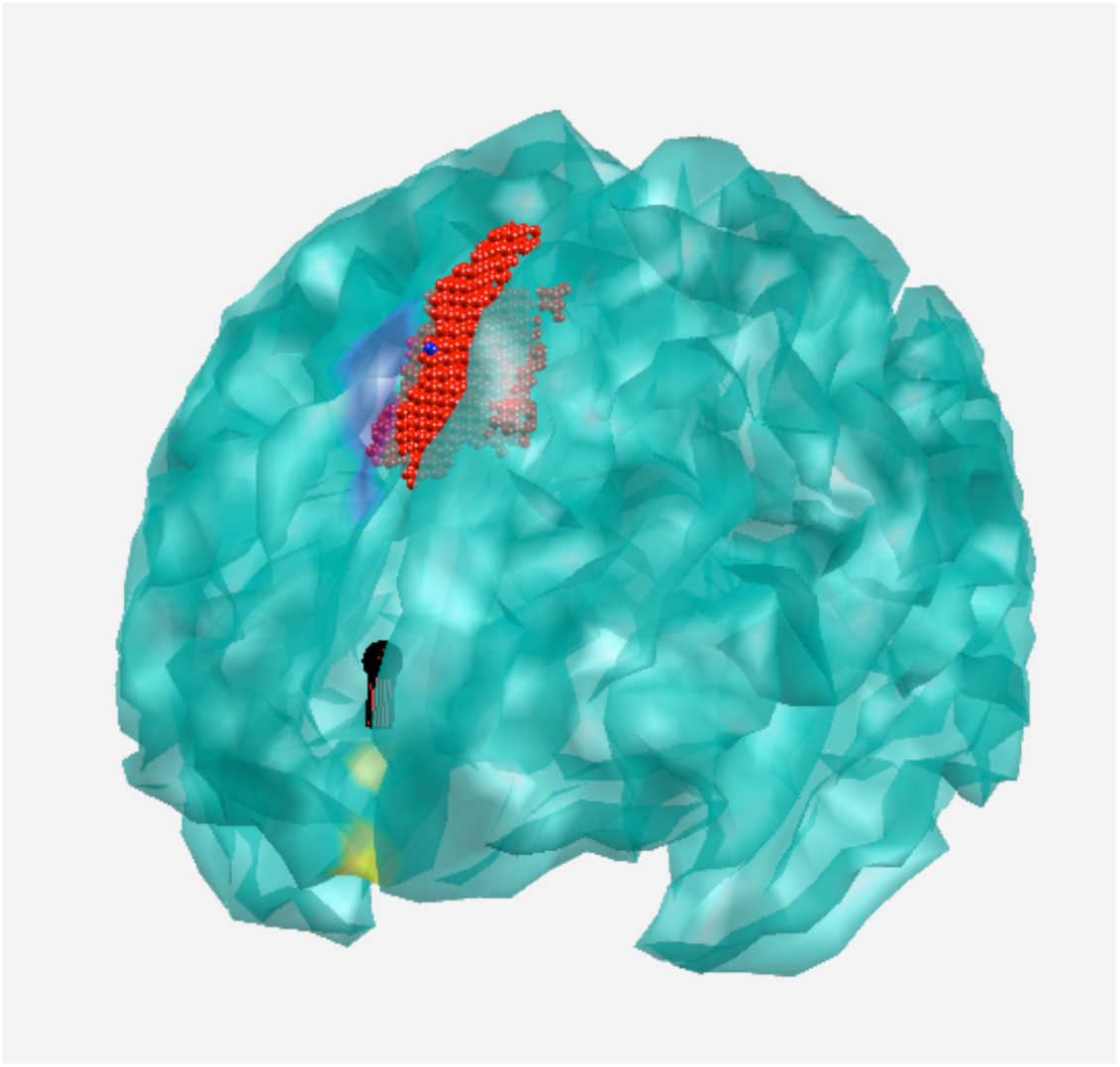
N450 Source Analysis. The ball marks the coordinates if the generator were a dimensionless point and the stem indicates the orientation of the positive side of the dipolar field. The yellow coloring indicates the implied cortical generator site if it were located in the positive direction of the field and the blue coloring indicates the implied cortical generator site if in the negative direction (either is a valid solution for these coordinates). The red marking is the nearest cluster [2 16 46] from the equivalent fMRI contrast (Table 4) to the negative implied cortical generator site.

**Table 4:**
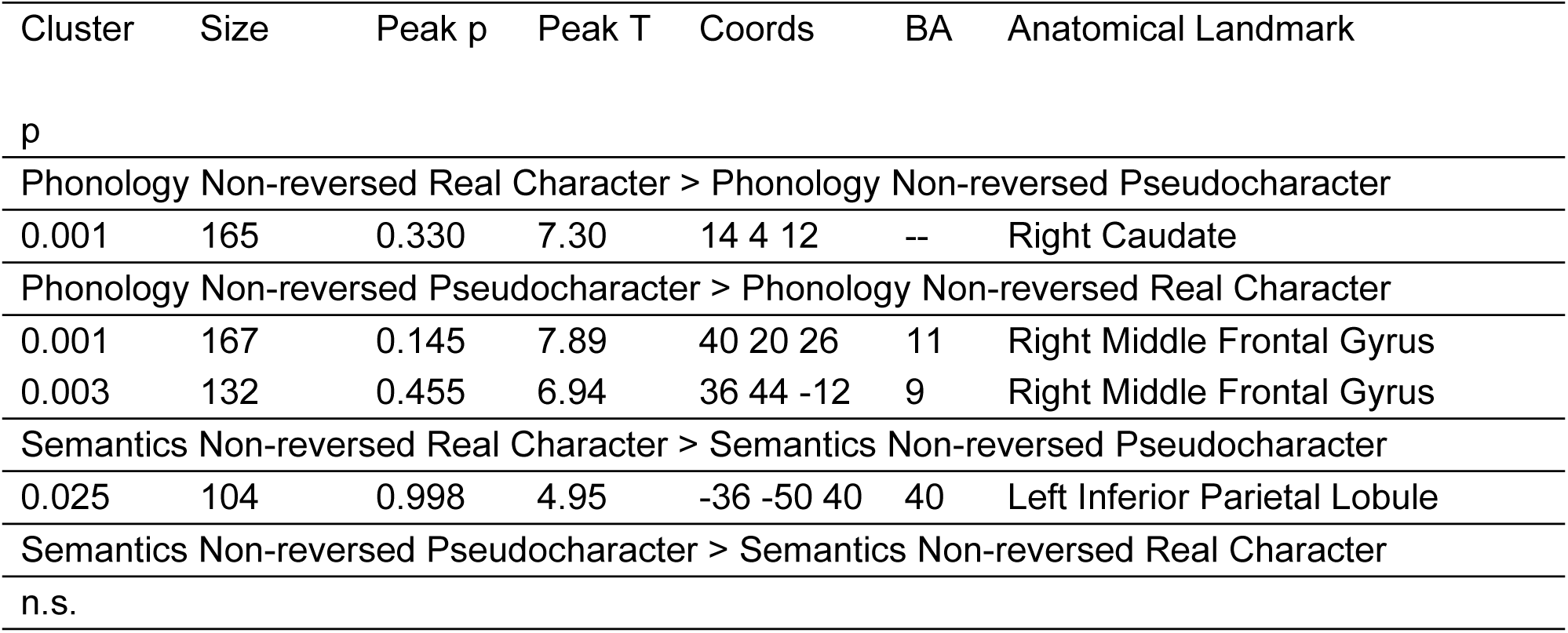
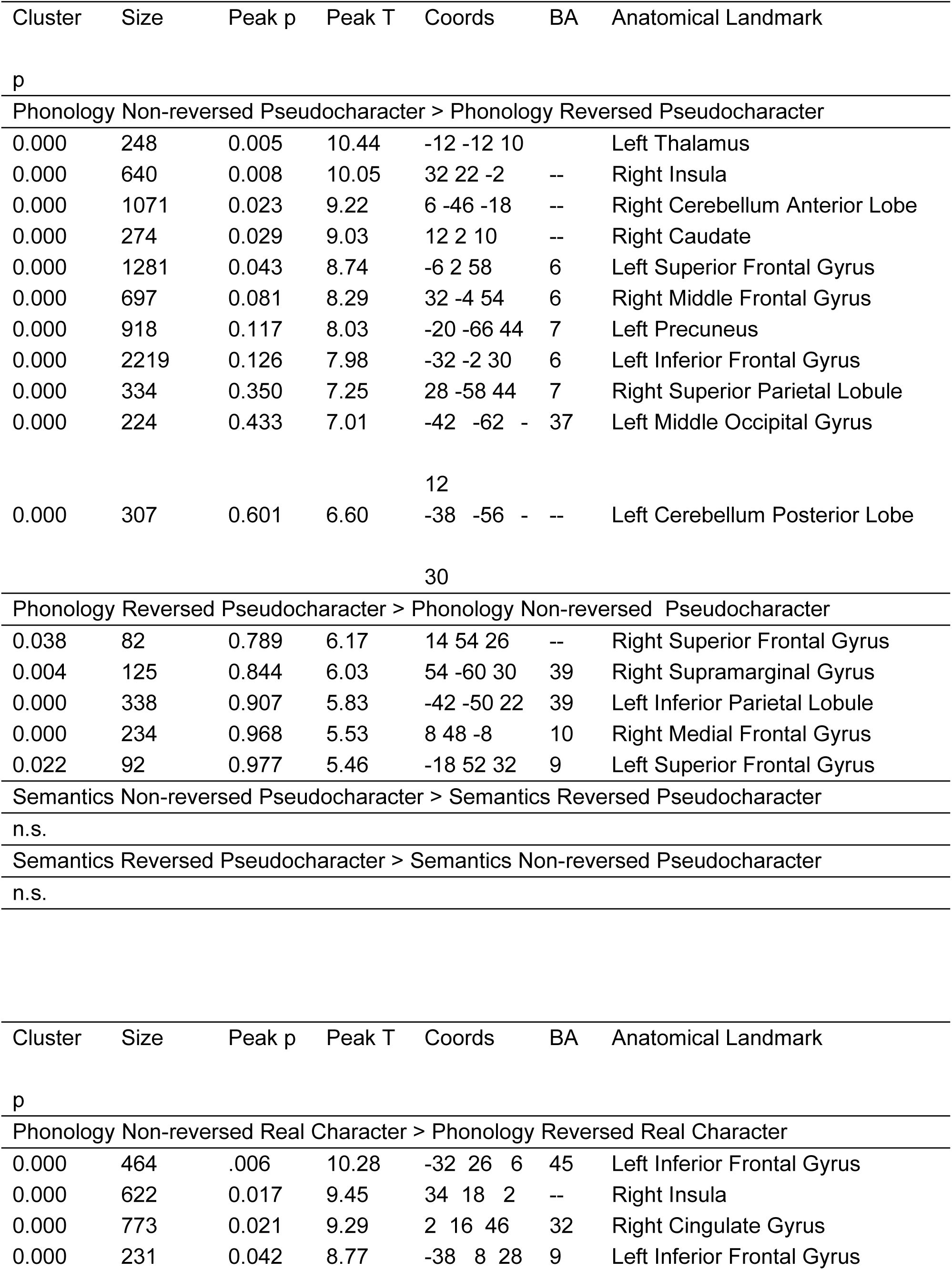
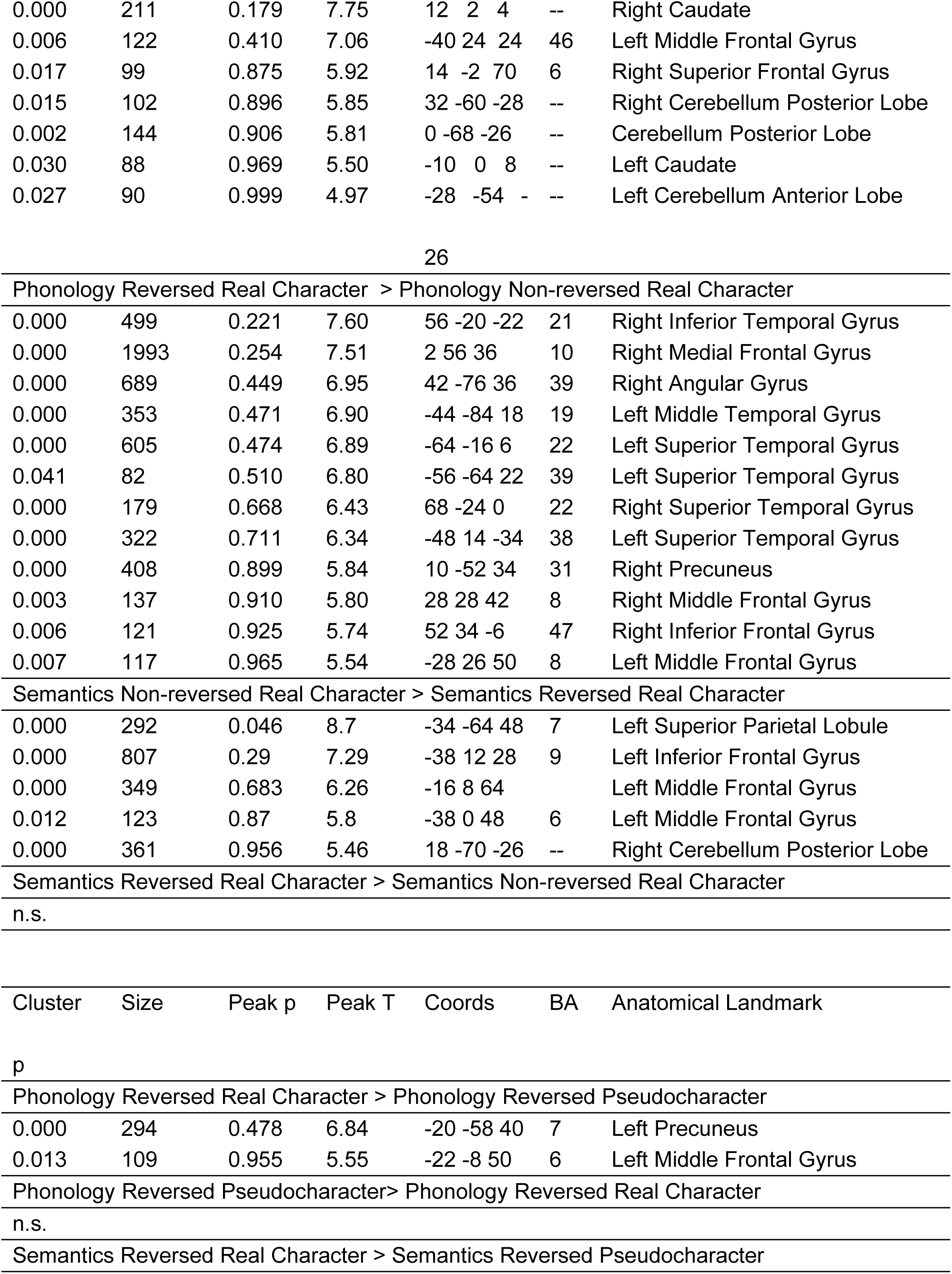
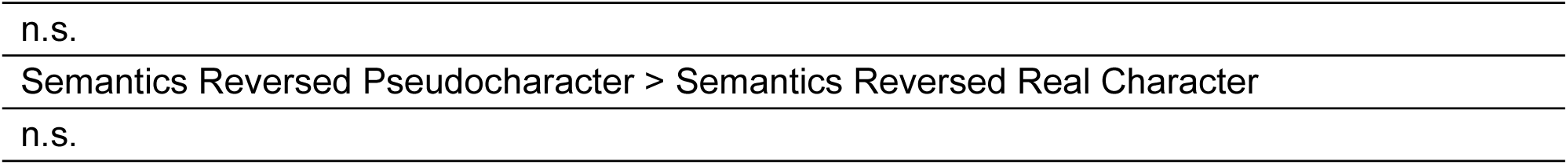
Planned Contrasts. Cluster p-values are FWE corrected. Size is number of voxels in the cluster. Peak p is FWE-corrected peak vertex p-value. Peak T is peak vertex value. Coordinates are MNI coordinates of the peak vertex. Vertex threshold set at p=0.001. BA is Brodmann Area.

A source analysis of the N170 using a difference wave (phonology reversed real character minus phonology non-reversed real character) at 180 ms resulted in point equivalent dipole MNI coordinates of [+/−38.06 −53.61 8.45], accounting for 79.39% of the variance (Figure 6). The implied negative cortical coordinates are [−45.6 −77.0 4.0] and [29.6 −66.3 −15.6], and the implied positive cortical coordinates are [−8.7 27.6 22.6] and [49.3 −32.6 46.0].

**Figure 6:**
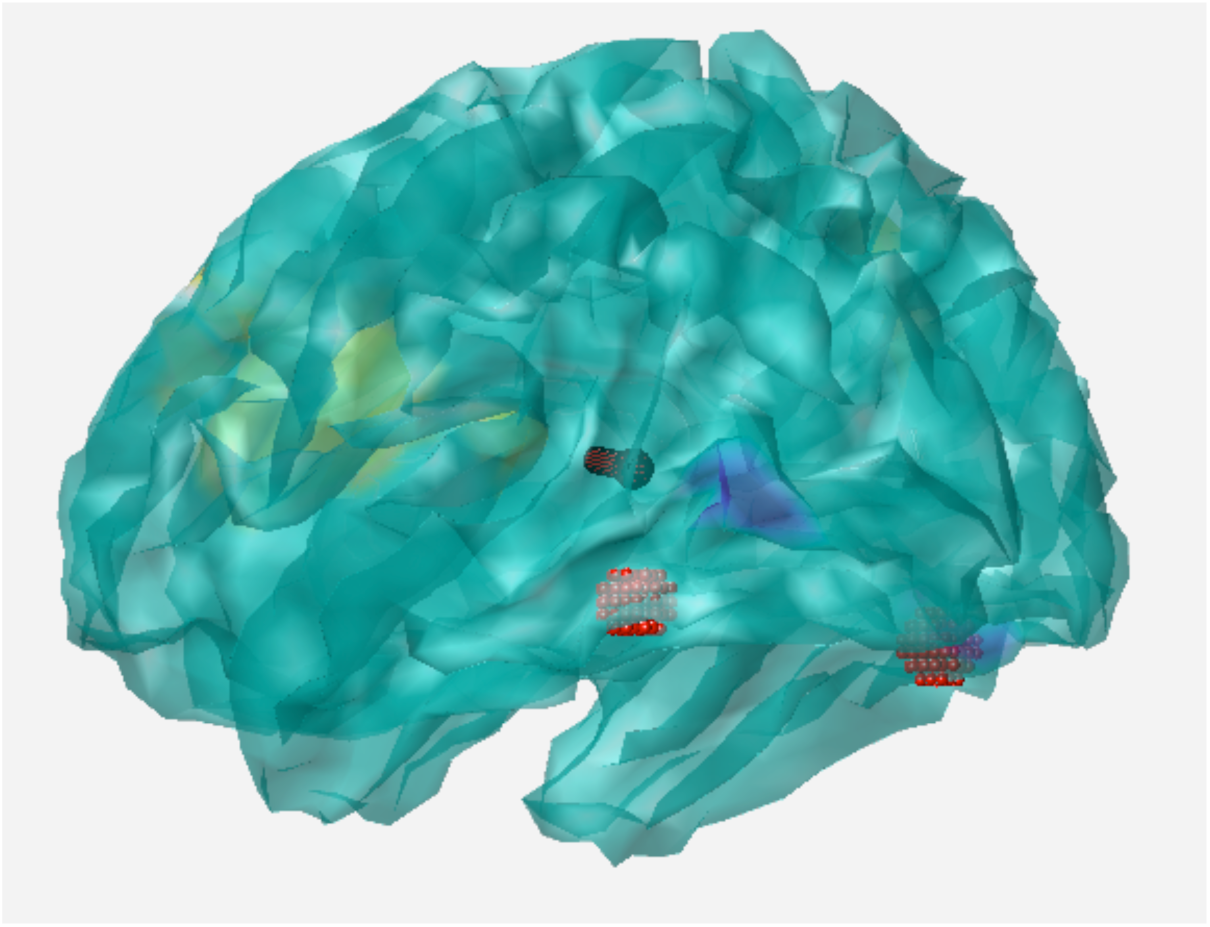
N170 Source Analysis. The ball marks the coordinates if the generator were a dimensionless point and the stem indicates the orientation of the positive side of the dipolar field. The yellow coloring indicates the implied cortical generator site if it were located in the positive direction of the field and the blue coloring indicates the implied cortical generator site if in the negative direction (either is a valid solution for these coordinates). The red markings are the VWFA ROI and its right hemisphere honologue.

### 2.3. ERP experiment discussion

The ERP data yielded clear N170 and N450 effects. The next step was to collect the fMRI data and then to co-register them with the source analyses. The same experiment scripts were used for the scanner sessions in order to maximize comparability between the sessions. Although ideally one might have wanted to collect EEG data simultaneously with the BOLD data, the high noise levels produced in the EEG data by the EPI sequences would have obscured the subtle N170 EEG effects. We also chose to use different participants as otherwise there would have been practice effects.

### 2.4. fMRI experiment material and methods

#### 2.4.1. Participants

Sixteen volunteers participated in the fMRI experiment in exchange for honoraria (4 male, 12 female; mean 23.6 years old, range 19-28), following a protocol approved by the University of Maryland Institutional Review Board. One was dropped due to equipment problems that resulted in incomplete data. All had normal or corrected-to-normal vision, were right handed, and were native Chinese mandarin speakers from the mainland and hence used to simplified characters. None reported any history of neurological, psychiatric, or learning disorders.

#### 2.4.2. Stimuli

The stimuli were the same as for the ERP sessions.

#### 2.4.3. Procedure

The procedure was the same as for the ERP sessions.

Stimulus presentation and recording of responses was implemented with E-Prime 2.0 (2.0.10.356), using an MRI compatible projection system (SilentVision SV-6011 LCD, Avotec Inc., Stuart, FL). Visual stimuli were projected onto a screen at the back of the magnet bore, viewed by participants through a mirror mounted on the MR headcoil. Responses were made via button-presses, using a fiber-optic button-box that registers latencies to the nearest ms.

#### 2.4.4. Data collection

The fMRI data were collected on a 3T Siemens MAGNETOM Trio Tim scanner with a 32-channel headcoil at the Maryland Neuroimaging Center at the University of Maryland, College Park. Functional image runs were obtained using a multiband echoplanar imaging (EPI) sequence (J. Xu et al., 2013) producing 60 oblique axial slices (TR/TE = 1250/39.4, flip angle = 90, field of view = 210 mm, matrix equal to 96×96, and isotropic voxels with a 2.2 × 2.2 × 2.2 mm resolution), covering the entire cerebrum and cerebellum. A high resolution, 3D anatomic image was acquired using a T-1 weighted (MP-RAGE) sequence (TR = 2100 ms, TE = 2.93 ms, TI = 1100 ms, flip angle = 12, 1 mm isotropic voxels, sagittal partitions) for the localization of functional activity.

#### 2.4.5. Data analysis

Data analysis was performed using Statistical Parametric Mapping (SPM12, http://www.fil.ion.ucl.ac.uk/spm). The images were time-corrected to the middle slice, realigned and unwarped, co-registered with the anatomical image and resliced into a 2×2×2 mm resolution, normalized by matching a mean echo planar imaging (EPI) image from each participant to the SPM canonical EPI East Asian brain template, and smoothed with a 6 mm full-width half-maximum kernel. ArtDetect-2015-10 (http://www.nitrc.org/projects/artifact_detect/) was used to detect bad reps with default settings and were nulled during analysis with rep-specific regressors. Initial statistical analyses utilized an event-related random effects approach with the informed basis set of the canonical hemodynamic response function, temporal derivative, and dispersion term, AR(1) correction for temporal autocorrelation (see Smith, Singh, & Balsters, 2007), default high-pass filter, and no global proportional scaling to avoid scaling artifacts (see Desjardins, Kiehl, & Liddle, 2001). In addition, six dimensions of head movement parameters plus a summary measure of head motion generated by ArtDetect were included in the model as nuisance covariates.

Clusterwise analyses were conducted using height thresholds set at p < 0.001, using FWE correction for multiple comparisons. Only the main term was used. Coordinates presented in this report are in MNI-space. Determination of anatomical features and Brodmann areas was performed using XJview (http://www.alivelearn.net/xjview). Results for peak voxel used if possible; otherwise, results for the largest gray matter portion of the cluster. Where closer inspection of putative Brodmann areas was required, the clusters were projected onto the Atlas (Talairach & Tournoux, 1988) using the Lancaster transform (Lancaster et al., 2007). MarsBar (Brett, Jean-Luc Anton, Valabregue, & Poline, 2002) was used for region-of interest (ROI) analysis and peri-stimulus temporal histogram (PSTH) figures using all three informed basis terms.

The VWFA ROI and the right hemisphere VWFA homologue ROI used the coordinates provided by the most similar prior fMRI study (Yang et al., 2011): [−42 −57 −12] and [42 −54 −15] as defined by their ICA. A 6 mm spherical ROI was drawn around each of these coordinates. The putative N450 ROI was computed by determining the Phonological Non-reversed Real Character versus Phonological Reversed Real Character cluster closest to the results of the source analysis and using it as the ROI. ROI analyses were performed by computing the percentage signal change (PSC) across the ROI and then using the mean PSC of the ROI as the dependent measure. While we also tried using the Target vs. False Strokes contrast as a localizer (Laurie Schwarz Glezer & Riesenhuber, 2013), the results were not sufficiently reliable at the single subject level. Instead, we used a group-level contrast as the localizer.

### 2.5. fMRI experiment results

#### 2.5.1. Behavior

Using trimmed means, overall semantic judgments (517 ms) were faster than phonology judgments (575 ms): T_WJt_/c(1.0,17.0)=29.28, p=0.00012, MSe= 4115. Also, overall responses to pseudocharacters (534 ms) were faster than to real characters (559 ms): T_WJt_/c(1.0,17.0)=17.80, p=0.00052, MSe= 1299. Additionally, overall responses to reversed characters (503 ms) were faster than to non-reversed characters (590 ms): T_WJt_/c(1.0,17.0)=203.95, p<0.00000001, MSe= 1318. These main effects were modulated by an interaction between Task and Lexical factors (T_WJt_/c(1.0,17.0)=7.77, p=0.015, MSe= 1389) such that the Task effect was more significant for real characters (T_WJt_/c(1.0,17.0)=51.87, p<0.00000001, MSe= 1961) than for pseudocharacters (T_WJt_/c(1.0,17.0)=8.35, p=0.013, MSe= 3543). There was also an interaction between the Task and Reversal factors (T_WJt_/c(1.0,17.0)=122.69, p<0.00000001, MSe= 634) such that the Task effect was significant for non-reversed characters (T_WJt_/c(1.0,17.0)=72.43, p<0.00000001, MSe= 2707) but not reversed characters. There was also an interaction between the Lexical and Reversal factors (T_WJt_/c(1.0,17.0)=36.83, p=0.000060, MSe= 1196) such that the Lexical effect was more significant for non-reversed characters (T_WJt_/c(1.0,17.0)=28.68, p=0.000060, MSe= 2283) than reversed characters (T_WJt_/c(1.0,17.0)=7.92, p=0.019, MSe= 211). Finally, there was a three-way interaction between all three factors (T_WJt_/c(1.0,17.0)=15.01, p=0.00084, MSe= 477) such that the Task by Lexical interaction was only significant for non-reversed characters: T_WJt_/c(1.0,17.0)=11.51, p=0.0030, MSe= 1543; this interaction in turn was more significant for the phonology judgement (T_WJt_/c(1.0,17.0)=22.85, p=0.00014, MSe= 3315) than for the semantic judgement (T_WJt_/c(1.0,17.0)=14.70, p=0.0025, MSe= 511).

The accuracy analyses yielded a similar pattern. Overall semantic judgments (100%) were more accurate than phonology judgments (98%): T_WJt_/c(1.0,17.0)=85.91, p<0.00000001, MSe= 0.00017. Also, overall responses to reversed characters (100%) were more accurate than to non-reversed characters (98%): T_WJt_/c(1.0,17.0)=54.01, p<0.00000001, MSe= 0.00032. There was also an interaction between Task and Lexical factors (T_WJt_/c(1.0,17.0)=40.25, p=0.000020, MSe= 0.000559) such that the Task effect was more significant for real characters (T_WJt_/c(1.0,17.0)=42.87, p<0.00000001, MSe= 0.001097) than pseudocharacters (T_WJt_/c(1.0,17.0)=15.78, p=0.0022, MSe= 0.00023). Finally, there was an interaction between Task and Reversal factors (T_WJt_/c(1.0,17.0)=56.90, p<0.00000001, MSe= 0.00024) such that the Task effect was significant for non-reversed characters (T_WJt_/c(1.0,17.0)=71.59, p<0.00000001, MSe= 0.00040) but not reversed characters.

#### 2.5.2. fMRI

For the localizer ROI, the closest significant cluster had a peak voxel at [−50 −58 −8] with a size of 376 voxels and cluster-level significance of p < 0.001. Closer inspection suggests it was more located in the posterior inferior temporal gyrus than the fusiform gyrus (Figure 7). There was a trend towards more activation for real characters versus pseudocharacters: TWJt/c(1.0,14.0)=3.78, p=0.07, MSe= 0.01. There was more activation for non-reversed versus reversed characters: TWJt/c(1.0,14.0)=6.33, p=0.026, MSe= 0.02. This effect was mediated by an interaction with task (TWJt/c(1.0,14.0)=7.93, p=0.017, MSe= 0.03) such that the reversal effect was only significant for the rhyming task: TWJt/c(1.0,14.0)=20.69, p=0.00024, MSe= 0.02. There was also a borderline significant three way interaction: TWJt/c(1.0,14.0)=4.40, p=0.06, MSe= 0.02. In the right hemisphere homologue [50 −58 −8] the three way interaction was significant (TWJt/c(1.0,14.0)=9.15, p=0.0098, MSe= 0.01) such that only for the rhyming task the lexical by reversal interaction was significant (TWJt/c(1.0,14.0)=8.75, p=0.012, MSe= 0.01), which in turn had the form of real characters producing more activation (TWJt/c(1.0,14.0)=6.81, p=0.024, MSe= 0.01) for reversed characters only.

**Figure 7:**
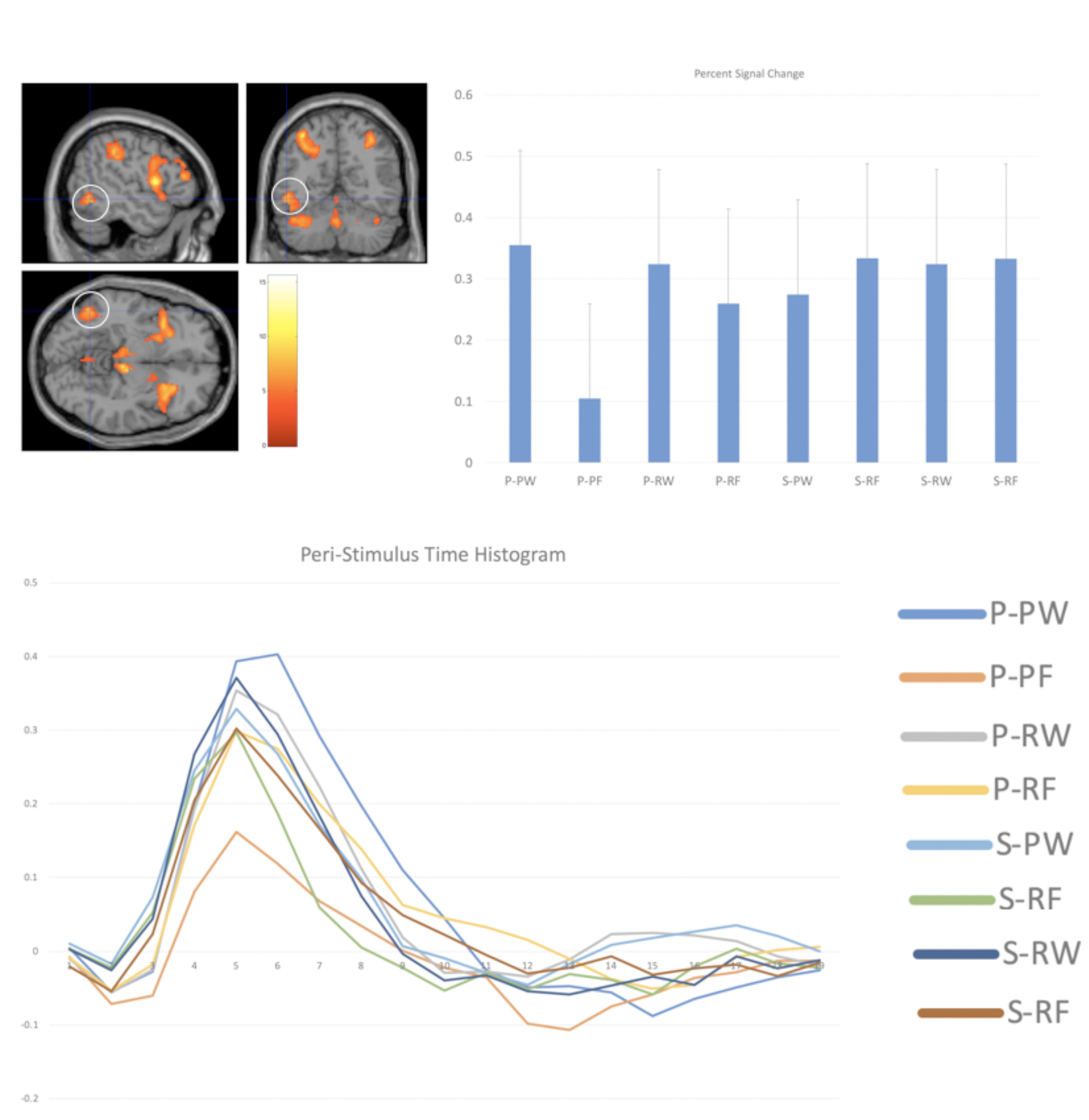
VWFA Localizer. The contrast was targets versus false strokes. The white circle marks the closest cluster to the VWFA. The error bars are the Loftus-Masson standard error (Loftus & Masson, 1994).

For the left hemisphere VWFA ROI (Figure 8) based on the prior study (Yang et al., 2011), centered on [−42 −57 −12], there was a significant effect of more activation for non-reversed characters: TWJt/c(1.0,14.0)=5.58, p=0.033, MSe=.001. This effect is moderated (TWJt/c(1.0,14.0)=8.92, p=0.018, MSe=.00078) such that it only occurs for pseudocharacters: TWJt/c(1.0,14.0)=9.35, p=0.011, MSe=.0013. There was also a borderline interaction of Task and Lexical (TWJt/c(1.0,14.0)=4.72, p=0.054, MSe=.001) that, if it were significant, would be due to a stronger phonology task effect for real words than pseudocharacters. Finally, for the right hemisphere VWFA ROI [42 −57 −12], there was an interaction between all three factors (TWJt/c(1.0,14.0)=5.12, p=0.043, MSe=.0022) but none of the follow-up tests were significant.

**Figure 8:**
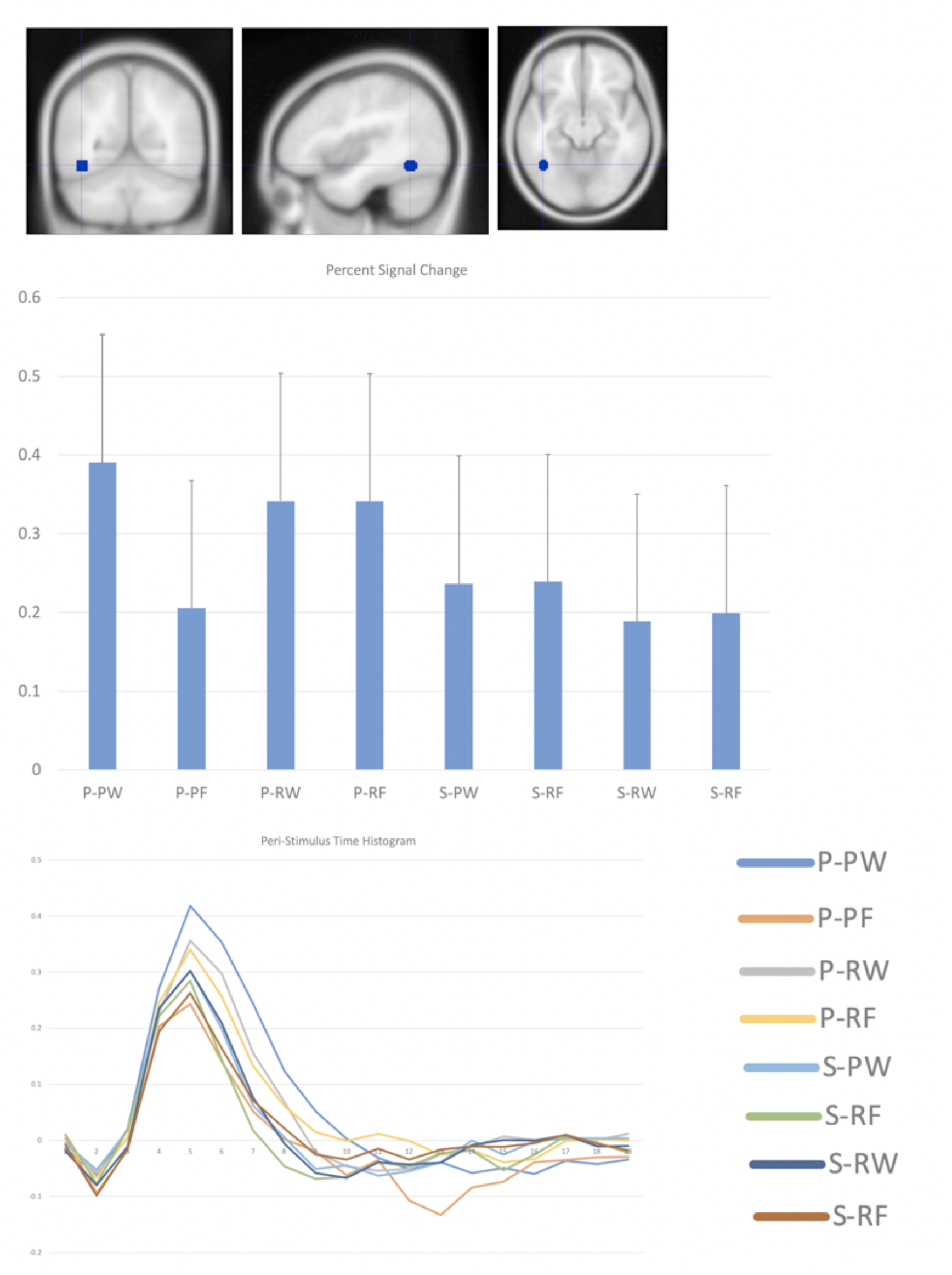
VWFA ROI. The blue circle marks the 6 mm radius ROI centered on the peak voxel of the VWFA region reported in a prior study (Yang et al., 2011). The error bars are the Loftus-Masson standard error (Loftus & Masson, 1994).

For the N450 ROI (Figure 9), the closest to the dorsal implied cortical generator site [7.9 31.6 50.4] was the cluster with a peak at [2 16 46]. Although xjVIEW labels this as being anterior cingulate BA 32, closer inspection suggests it is better described as being supplementary motor cortex BA 6. In this ROI, there was more activation for non-reversed characters: T_WJt_/c(1.0,14.0)=61.46, p<0.00000001, MSe= .0012. This effect was moderated by a Task interaction (T_WJt_/c(1.0,14.0)=22.89, p=0.0012, MSe= 0021) such that this effect of reversal was only significant for the phonological judgements: T_WJt_/c(1.0,14.0)=126.43, p=0.000020, MSe= .00096. This interaction too was moderated by an interaction with the Lexical factor (TWJt/c(1.0,14.0)=8.05, p=0.018, MSe= .0015), such that it was more significant for pseudocharacters (T_WJt_/c(1.0,14.0)=22.71, p=0.0010, MSe= .0024) than real characters (T_WJt_/c(1.0,14.0)=4.86, p=0.047, MSe= .0012).

**Figure 9:**
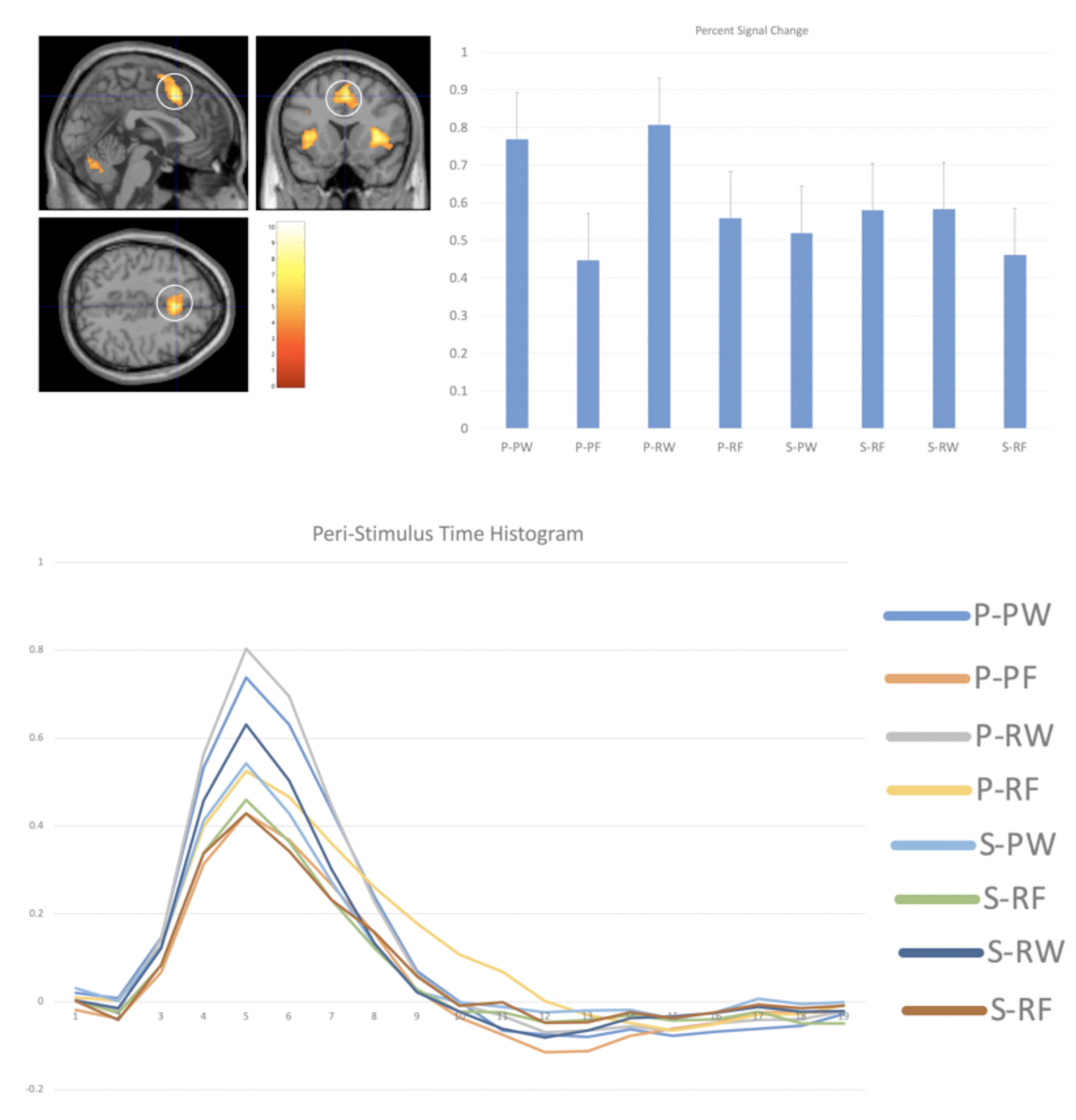
N450 ROI Results. The contrast was phonology un-reversed real characters versus phonology reversed real characters. The white circle marks the closest cluster to the implied N450 cortical generator, which was used as the ROI. The error bars are the Loftus-Masson standard error (Loftus & Masson, 1994).

## 3. General discussion

The ERP and the fMRI data yielded results most consistent with the order-sensitive Lexicon hypothesis: (RW=PW)>(RF=PF). The N170 and N450 effects were observed in the ERP data and the VWFA effects were observed in the fMRI data. This conclusion comes with a number of caveats and additional observations.

The initial attempt to establish a VWFA ROI via a group-level localizer was not entirely successful. We contrasted the target condition with the false strokes condition, reasoning that since neither condition was being included in the analyses, we would be safe from double-dipping (Kriegeskorte, Simmons, Bellgowan, & Baker, 2009). While we did obtain a plausible looking cluster, it was mostly located in the adjoining posterior inferior temporal gyrus and middle occipital gyrus. While not unusual (see Mechelli et al., 2003; Price et al., 1996), this is part of the Language Formulation Area (Nielsen, 1946) that has been proposed to subserve the somewhat different function of indexed phonology (Dien et al., 2013). Indeed, there was greater activation for the non-reversed characters in the rhyming task only, as one would for a region mediating phonological access; furthermore, there was a borderline effect of more activation for real characters versus pseudocharacters, as one would also expect from a region mediating indexed phonology.

Reasoning that our informal localizer had provided insufficient statistical power compared to more optimized methods (Laurie Schwarz Glezer & Riesenhuber, 2013), we opted instead to utilize an ROI based on the prior study (Yang et al., 2011) most closely resembling the present study. Here, the critical effect was that non-reversed pseudocharacters (PW) produced more activation than reversed ones (PF). Although the order-sensitive Lexicon hypothesis would have predicted the same pattern for real characters, this omission could have been due to insufficient statistical power.

Turning to the N170, we find that the source analysis was in general consistent with it emanating from the VWFA region, within the limited spatial resolution of such analyses. If we accept this proposition, we find that it also showed a reversal effect and it did not distinguish between the real and pseudocharacters, supporting the notion that the fMRI data simply missed the effect for real characters. Interestingly, the effect the N170 showed was not one of amplitude but rather a prolongation.

While our finding of a prolonged N170 differs from that of a prior study (Lin et al., 2011) that also contrasted pseudocharacters and false characters and reported a smaller amplitude to the reversed characters, it is likely that this difference is because they used an implicit task of judging the font color and so participants did not need to identify the characters.

Although one might expect such a prolonged ERP activation to correspond to increased BOLD signal, instead it was the reverse. It may be that discrepancy was due to the differing time sensitivities of the two methods, with the ERP revealing a subtle change in the initial phasic burst of activity to reversed characters that then subsided as the system rejected the false word form, while the fMRI responded to a small but steady top-down tonic activation for the rest of the trial, as hypothesized by the Interactive hypothesis. This notion of extended processing in the VWFA is also consistent with observation of increased activity for words with lower frequency and consistency (Graves, Desai, Humphries, Seidenberg, & Binder, 2010; Kronbichler et al., 2004).

While the N170 data was overall most consistent with order-sensitivity (as it responded to reversal), it was intriguing that the nature of its response was a prolongation of the initial phasic burst. It may be that the better description of it is semi-order-sensitive, in that it can detect reversal but this manipulation is sufficiently challenging for the system that it needs some additional processing time to come to a resolution. The hypothesis (Frankish & Turner, 2007) that the orthographic pathway is less order-sensitive than the phonological pathway is therefore still viable.

The N450 data provided additional insights into the experiment. Although it had not been previously established that this kind of rhyming task (responding to a previously instructed target sound) could elicit an N450 rhyming effect (the usual design involves sequential pairs of stimuli), the present experiment displays a robust N450 effect. Furthermore, it confirms that pseudowords do activate sounds in the minds of the participants (Lee et al., 2010). It extends these findings by also indicating that reversed radical characters do not activate sounds (or at least to a lesser degree).

While not necessary for interpreting the present results, the co-registered data suggests for the first time a generator site for the N450 effect, namely the supplementary motor area or SMA. Other fMRI rhyming studies have also reported effects in the SMA (Donald J. Bolger, Hornickel, Cone, Burman, & Booth, 2008; Kim et al., 2016; L. Liu et al., 2009). This site is consistent with it being produced by subvocalization. The fact that the participants apparently felt the need to subvocalize to perform the rhyming task, and at such a long latency, raises the interesting question as to what degree phonological decoding is completed prior to semantic access (Perez-Abalo et al., 1994).

One limitation of this study is that the behavioral data shows that the rhyming task was more difficult than the semantic task, so any task effects are confounded with difficulty. Fortunately, the conclusions of this study do not rely on the task effects.

Another limitation of this study is that while it sufficed to test the level of order sensitivity posited by the LCD model (Dehaene et al., 2005), a more complex design would be required to test that of competing models. For example, the SERIOL model, which implements open bigrams (Grainger & Whitney, 2004) which are like local bigrams but with an activation gradient that extends across the entire word (C. Whitney, 2001; C. Whitney & Berndt, 1999), includes an additional component that gives the first and last letters special status (Carol Whitney & Cornelissen, 2008), recognizing studies that suggest they play a greater role in word recognition (Guerrera & Forster, 2008; T. R. Jordan & Patching, 2003; Timothy R. Jordan, 1995; Schoonbaert & Grainger, 2004). From this standpoint, the radicals in the present study are equivalent to such exterior letters. The nature of order sensitivity was not the central subject of this present study and so it does not affect its primary conclusions, but does point towards the need for further studies.

## 4. Conclusions

In summary, this study has applied both ERP and fMRI methods and the special properties of the Chinese script to test three hypotheses of the VWFA and have concluded that the Lexicon hypothesis is best supported. Furthermore, we have distinguished between order-sensitive and order-insensitive versions of these models and have concluded that the data supports the order-sensitive version, although it may be that the better description would be semi-order-sensitive. As a bonus, we also provide evidence that the N450 rhyming effect arises from the supplementary motor area, which may help inform future studies using this effect.

## Acknowledgements

This work was supported by the Fundamental Research Funds for the Central Universities (3102016ZY042). The funding source had no involvement at any step beyond the granting of the funds. Thanks to Tianyin Ouyang for her invaluable assistance.

